# Age-Dependent Maturation and Rejuvenation of the Neural 3D Chromatin Interactome in Enriched Environments

**DOI:** 10.1101/2024.10.22.619383

**Authors:** Javier Gancedo-Verdejo, Rocío G. Urdinguio, Juan Ramón Tejedor, Raúl F. Pérez, Alfonso Peñarroya, Pablo Santamarina-Ojeda, Annalisa Roberti, Jennifer M. Kefauver, Carlota Álvarez-Díaz, Mar Rodríguez-Santamaría, José Luis Trejo, Laureano Tomás-Daza, Llorenç Rovirosa, Biola M. Javierre, Agustín F. Fernández, Mario F. Fraga

## Abstract

Aging is a multifactorial biological process resulting in physiological and cellular decline. However, our understanding of age-related changes in 3D genome organization and the effect of external interventions on this process, remains limited. Here we describe alterations in the landscape of the 3D chromatin interactome upon aging, utilizing the low input Promoter Capture Hi-C (liCHi-C) technique with hippocampal neurons. We integrated liCHi-C data with RNA-seq data to identify functional implications. Furthermore, we assessed the effect of exposure to environmental enrichment (EE). Remarkably, our results demonstrated an age-dependent modulation of promoter interactions and expression with EE, with aging-like changes induced in young mice upon EE, likely associated with early brain maturation; while age-related alterations were reverted in old mice, leading to a partial rejuvenation of aged mouse hippocampi. These findings revealed a dynamic behaviour of the neuronal 3D chromatin structure over time, which can be modulated by external interventions.

## INTRODUCTION

Aging is a multifactorial biological process characterized by a progressive decline in physiological and cellular functions as a result of molecular alterations accumulated throughout lifespan^1,2^. This complex process is associated with an increased risk of age-related disorders, such as cancer, cardiovascular diseases, or neurodegenerative disorders^1^. In the last years, several molecular mechanisms of aging have been described, with epigenetic alterations playing a significant role in this process, as well as in aging-associated diseases^1,3–5^. The effect of aging on various brain regions has been extensively studied at different molecular levels, and especially in the hippocampus, which is crucial for cognition and memory, and has active neurogenesis in adults^6,7^. Epigenetic and chromatin alterations in hippocampal cells during aging have been related to cognitive decline^6,8^ and associated with neurodegenerative diseases^9^. Moreover, these epigenetic mechanisms can be modulated by environmental factors, ultimately impacting gene expression patterns^10–12^. Nowadays, epigenetic regulation is known to play a critical role in cognition^13^ and memory^14^, neurogenesis^15,16^, neuronal plasticity^17^ and neurodegeneration^18^, many of which are affected by aging, stress, and environmental factors^15,19^.

Some of the mechanisms involved in epigenetic regulation are those related to chromatin conformation. Chromatin presents a complex three-dimensional (3D) organization that plays an important role in the regulation of gene expression^20,21^ and the determination of cell identity^22^. The regulatory activity of certain genomic elements, such as enhancers, is essentially controlled by the chromosomal structure. Enhancers, which are modulators of gene expression often located at a long linear distance from the gene, exert their regulatory function through spatial proximity to target gene promoters. These physical interactions are established by the 3D folding of chromatin into loops^23–26^. There are two types of chromatin loops: structural loops and functional loops, the latter being determined by those promoter-enhancer interactions^27^. Furthermore, cell type-specific expression patterns are determined by the combined effects of proximal and distal regulatory elements^20^.

Chromatin conformation capture (3C) technologies have been essential for the study of chromatin organization. Among them, the Hi-C technique allows for the identification of genome-wide interactions^27^ and facilitates the exploration of the 3D genome at different scales, including both chromatin interactions in *cis* (within chromosome) and in *trans* (between chromosomes)^23,28^. Hi-C techniques have provided valuable insights into mechanisms underlying cognition and memory^29–31^, brain development^32,33^, neurogenesis^34^, as well as neuropsychiatric and neurodegenerative disorders^35,36^. Furthermore, variants of the Hi-C technique have been performed to explore cerebellar aging^37^. However, due to the high cost and low resolution of the Hi-C technique, Capture Hi-C (CHi-C) approaches have been developed to enrich the sample library in specific genomic regions^38^. One such method is Promoter Capture Hi-C (PCHi-C), which enriches the library for promoter interactions using probes that hybridize with promoter fragments^39^. Among other findings, PCHi-C has provided insights into the mechanisms underlying neurodevelopmental diseases^40^ and neuropsychiatric disorders^41^. However, the characterization of aging-related alterations in promoter interactions using PCHi-C approaches remains limited.

Chromosomal organization varies greatly across different cell types, resulting in highly cell-type-specific chromatin interactions^33,42^. There is a significant cellular heterogeneity in the brain, being neurons and glial cells the predominant categories^43^. For that reason, different strategies have been employed to mitigate cellular heterogeneity for Hi-C analysis in the brain, including cell sorting^33,44^. However, the starting number of cells is a limitation of PCHi-C, which requires millions of cells; the low input Capture Hi-C (liCHi-C) method is an alternative suited for the analysis of very low cell numbers^45^.

Before this study, the age-dependent effect of the environment on the neuronal 3D chromatin interactome was unknow. Therefore, here we aimed to characterize the 3D chromatin interactome in hippocampal neurons of young and old mice and its association with gene expression to identify functional alterations that occur with age. Furthermore, we investigated the potential of external factors to modify chromatin contacts and gene expression. To achieve this, we exposed young and old mice to environmental enrichment (EE), using an established cognitive and physical enrichment design^30,46–48^, and we focused on the dorsal hippocampus, due to its involvement in learning and memory, as well as cognitive decline with aging^6^. Thus, with our approach we have characterized alterations in promoter interaction associated with gene expression changes in the context of aging. In addition, we observed an age-dependent impact of EE, where some age-related alterations are boosted by EE in young mice, while others are partially reversed in aged hippocampi.

## RESULTS

### 3D Chromatin Interactome changes with age

To characterize the three-dimensional (3D) chromatin structure in mouse dorsal hippocampal neurons and investigate how is it affected by age, we used a Hi-C approach. First we purified neuronal nuclei from young and old mouse hippocampi using the well-established intracellular marker NeuN^49^, and subsequently we performed low input Promoter Capture Hi-C (liCHi-C) (Figure 1A).

**Figure 1.**
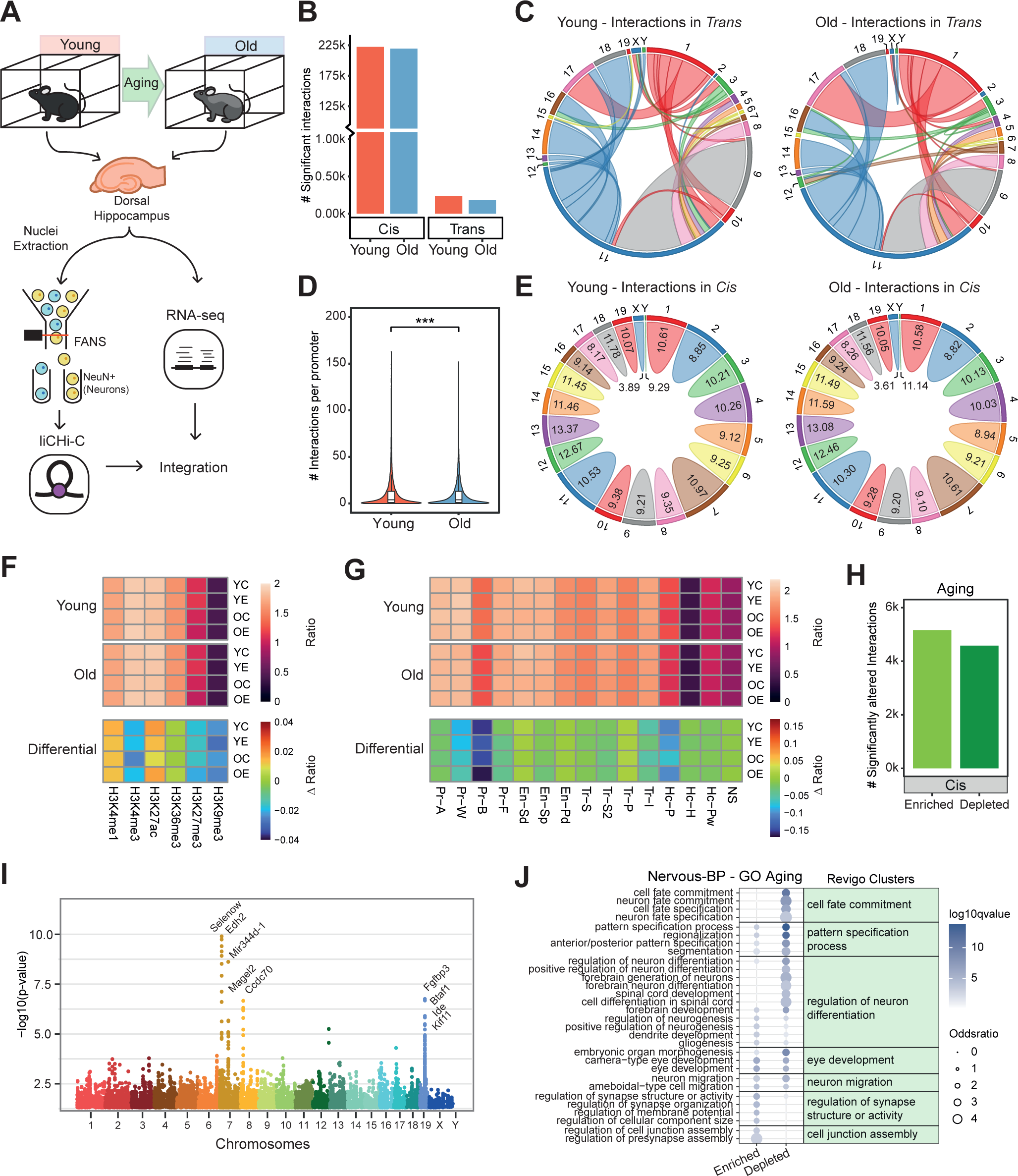
Promoter Interactions of young and old mice hippocampal neurons and their change with aging. **A**. Schematic of the study design for young and old mice conditions, using low input Promoter Capture Hi-C (liCHi-C) and RNA-seq analysis and performing integrative analysis. **B**. Bar plot illustrating the total number of CHiCAGO significant interactions in *trans* and in *cis* for young and old mice. 1000 (k). **C**. Chord plots representing the *trans* interaction proportion of each mice chromosome (outside number) for young mice (left) and old mice (right). **D**. Violin plots showing the number of interactions per promoter. Merged samples for young and old conditions were represented. Wilcoxon signed-sum test for paired samples was used to compare groups (p-value = 1.331e-07). ***p-value < 0.001. **E**. Chord plots representing the *cis* interaction of each mice chromosome (outside number) for young mice (left) and old mice (right). The width of each chord represents the proportion of *cis* interaction of each chromosome, and the inside numbers indicate the number of *cis* interactions per captured promoter in each chromosome. **F, G**. Heatmaps representing histone mark (**F**) and chromatin states (**G**) enrichment analysis of significant interactions in *cis* for young and old mice. Six common histone posttranslational modifications, and the 15 chromatin states (see Methods section) defined by histone marks, from four mouse hippocampus datasets (the four experimental conditions: YC, YE, OC and OE) of a previous study^47^ were used in the enrichment analysis. First color scale (on top) represents the magnitude of enrichment (ratio), and the second color scale (on bottom) reflects the differential ratio (Δ ratio) with aging (old vs young), calculated subtracting the young mice values from the old mice values. **H**. Bar plot illustrating the total number of Chicdiff interactions in *cis* that change significantly (p-value < 0.05) with aging, distinguishing between enriched and depleted interactions. 1000 (k). **I**. Manhattan plot of differential interactions in *cis* with aging (Chicdiff p-value < 0.05). **J**. Bubble plot represents Gene Ontology (GO) terms enrichment based on Nervous system-associated Biological Processes (Nervous-BP) for promoter genes with significantly enriched and depleted interactions (p-value < 0.05) with aging. Bubble color intensity represents the statistical significance (-Log_10_ q-value) and dot size reflects odds ratio. Top significant GO terms were grouped following REVIGO clustering, which are named with the most representative term of the group.

For liCHi-C, we isolated neuronal nuclei from young and old mouse brains using fluorescence-activated nuclei sorting (FANS) (Figure S1A). Mature NeuN^+^ neurons comprised approximately 60% of the hippocampal population (Figure S1B). Three biological replicates were performed, each using 1 million neuronal nuclei pooled from 3 individual mice. liCHi-C libraries were then prepared and deep-sequenced. Paired-end reads were processed, mapped, and filtered using HiCUP pipeline^50^. The percentage of valid reads (mean = 64.336%; SD = 2.416%), the capture efficiency (mean = 60.328%; SD = 3.251%) and the proportion of interactions in *cis* and in *trans* (*trans* mean = 14.882%; SD = 0.362%) revealed no significant differences between replicates of young and old mice (Figures S1C, S1D and Table S1), and demonstrated high quality of liCHi-C libraries. Subsequently, the CHiCAGO pipeline^51^ was used for significant interaction calling (CHiCAGO score ≥ 5), detecting more than 200,000 significant contacts in merged samples (mean = 221,560; SD = 1,966). The proportion of promoter-promoter interactions remained consistent among replicates, representing around 14.4% of all interactions compared to promoter-to-non-promoter interactions (Figures S1E, S1F and Table S1).

From CHiCAGO interaction calling, we identified 239 CHiCAGO significant contacts in *trans* for young mice, while there were 181 for old mice. For instance, interactions between chromosomes 11 and 9 seemed to be reduced in older mice (Figures 1B, 1C and S1G). In contrast, both young and old mice displayed over 200,000 CHiCAGO significant contacts in *cis* (222,484 in young and 219,779 in old) (Figure 1B). At a global level, the number of *cis* contacts per promoter was significantly lower in older mice than younger mice (Wilcoxon p-value < 0.001), and chromosome 13 harboured the highest number of interactions in *cis* per captured promoter in each condition (Figures 1D, 1E). Moreover, distance distribution of *cis* interactions differed between age groups, as young mice exhibited a significantly greater median distance (263 kb) between interacting regions, compared to old mice (254 kb) (Wilcoxon p-value < 0.001) (Figure S1H and Table S1).

Due to their abundance and potential functional implications, we focused on *cis* interactions, performing an integration with histone modifications / chromatin states obtained from mouse hippocampus^47^. We observed an enrichment of activating histone modifications (e.g. H3K4me3, H3K27ac) in young and old mice. However, a slight decrease in H3K4me3 mark as well as in repressive marks (e.g. H3K9me3) was observed with aging (Figure 1F). Additionally, *cis* interactions were enriched for chromatin states related to promoters and enhancers, but not heterochromatin states. Comparing the two age groups, we found a decrease in the enrichment of some chromatin states with aging. These included bivalent and weak promoters, and polycomb-associated heterochromatin (Figure 1G). We had previously described a reduction of bivalent promoter states with aging^47^, while here we observed a reduction of interactions with bivalent promoter states, supporting the idea that, at chromatin accessibility and organization level, bivalent promoters are highly affected by the aging process^52^.

We next performed a differential analysis using the Chicdiff pipeline^53^, which identifies significant changes in promoter interactions between conditions by pooling interaction counts from contiguous fragments to improve the statistical power. This analysis revealed 9,740 interactions that changed with age (p-value < 0.05), 5,162 interactions showing significant enrichment and 4,578 showing significant depletion (Figure 1H). When we analyzed their association with common histone marks, in promoter regions, an increase of active histone marks was observed, specially H3K27ac and H3K4me3. However, repressive marks (e.g. H3K27me3 or H3K9me3) did not suffer significant changes in promoters (Figures S2A-F). Furthermore, in the interacting regions of significantly changed contacts, similar, though slighter, observations were detected (Figures S2A-F).

Regarding their genomic location, some of the interactions that were most significantly altered with age clustered within specific regions, such as those in chromosome 7 (involving *Selenow* and *Edh2* promoters) or in chromosome 19 (involving *Fgfbp3*, *Btaf1*, *Ide* and *Kif11* promoters) (Figure 1I). Furthermore, gene ontology (GO) analysis of promoters with enriched interactions in old mice revealed an association with synapse organization and transcription initiation processes (Figures 1J, S1I and Table S2), suggesting a possible role in synaptic plasticity during aging^54^. Conversely, genes with depleted interactions in old mice were mainly linked to functions like cell fate commitment, pattern specification, and neuron differentiation (Figures 1J, S1I and Table S2), indicating a potential role in cell identity^41^. These results suggest that age-related changes in chromatin interactions are not random but potentially influence the regulation of specific biological pathways.

### Transcriptomic analysis reveals age-related changes and potential chromatin interactome link

To assess the impact of aging on the transcriptome and its potential connection to the chromatin interactome, we performed RNA sequencing (RNA-seq) on dorsal hippocampus tissue from mice (Table S3). Since sorting NeuN^+^ cells from the dorsal hippocampus requires membrane permeabilization for labelling this intracellular marker, resulting in the loss of mRNA molecules, RNA-seq was performed using bulk hippocampal cells, following previous experimental designs^30^. Subsequent integration with liCHi-C data allowed us to describe changes in promoter interactions with a possible influence on gene expression regulation (Figure 1A).

PCA of gene expression shows a segregation of samples by age (Figure 2A). Differential expression analysis identified genes whose expression levels were significantly altered with aging. Specifically, we found 1,116 genes significantly upregulated, including immune-related genes like *C4b* and *Ighm*. Conversely, 954 genes were significantly downregulated with age, including *Nrep*, a gene involved in axon regeneration (Figures 2B, 2C). To explore the functional implications of these age-related gene expression changes, we performed a Gene Set Enrichment Analysis (GSEA)^55^ on expression data. This analysis identified 56 gene sets significantly upregulated with aging. Many of these upregulated gene sets were associated with ribosomal regulation and translation processes (Figure 2D and Table S4). Conversely, 34 gene sets exhibited significant downregulation in aged mice. These downregulated sets included those related to G protein-coupled receptors (GPCRs), brain markers, and neurogenesis (Figure 2D and Table S4).

**Figure 2.**
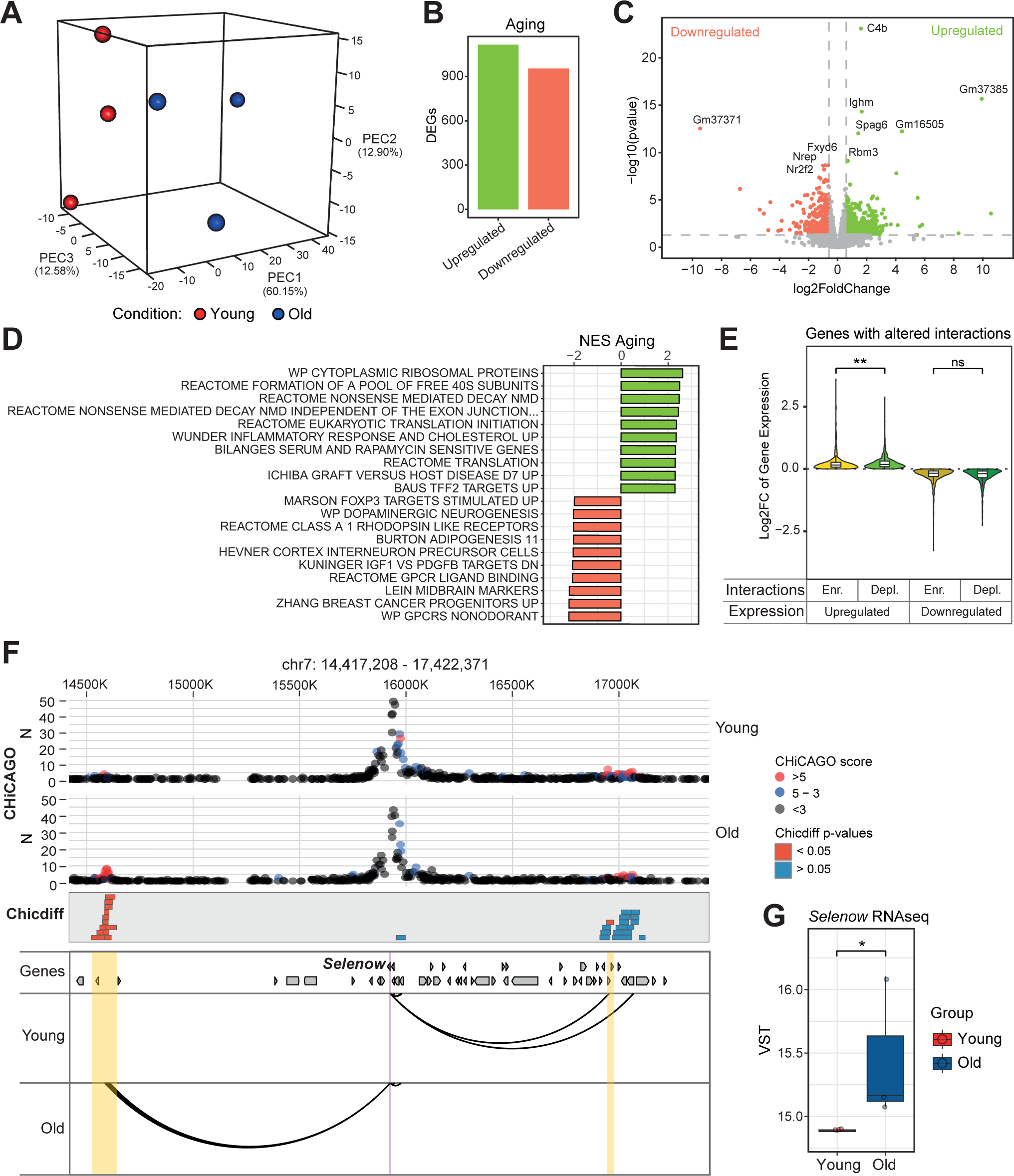
Gene expression alterations with aging and integration with liCHi-C data. **A.** Principal Component Analysis (PCA) for all expressed genes of RNA-seq replicates in young and old conditions. The proportion of variance explained for the PC1, PC2 and PC3 components is indicated. **B**. Bar plot illustrating the number of Differentially Expressed Genes (DEGs) with aging (old vs young), distinguishing between upregulated and downregulated genes (p-value < 0.05). **C**. Volcano plot showing the significance and the fold change of the upregulated genes (green dots) and downregulated genes (red dots) with aging. **D**. Barplots reflecting the Normalized Enrichment Score (NES) of GSEA gene sets from M2 collection (MSigDB). Gene sets NES were calculated for aging (old vs young). Top significantly enriched (green) and depleted (red) gene sets are shown (FDR q-value < 0.05). Extended lists of significant gene sets are available in Table S4. **E.** Violin plots showing the Log_2_ Fold Change of genes with altered interactions, splitting between genes upregulated or downregulated with aging, and comparing those with enriched or depleted interactions with aging. Significance calculated with Wilcoxon rank-sum test. **F**. *Selenow* promoter-centered interactions according to liCHi-C data in young and old conditions. Top: Dots represent counts of contacts, highlighting the CHiCAGO significant ones in red. Rectangles indicate significantly altered interactions (p-value) between conditions based on Chicdiff analysis. Bottom: Arcs represent CHiCAGO significant contacts. Purple shade depicts *Selenow* gene promoter and yellow shade depicts regions with significantly altered interactions. Arrows symbolize gene placement and orientation along the genomic window. **G**. Boxplots indicating the RNA-seq expression measurements (VST normalized data) for the *Selenow* gene across young and old mice. Dots denote individual mouse. ^ns^p-value > 0.05, *p-value < 0.05, **p-value < 0.01.

We next explored the relationship between chromatin interactions and gene expression by integrating liCHi-C and RNA-seq data. Despite observing a higher increase of expression in genes with significantly enriched interactions (Figure 2E), a clear and consistent pattern across different interaction and expression combinations went unnoticed (Figures 2E, S3A and Table S5). This finding supports the idea that the direction of the changes in interactions (enriched or depleted) might not directly translate to predictable changes in gene expression (upregulation or downregulation)^40^, or that not all chromatin interactions have a gene transcription regulatory role. This highlights the complexity of chromatin interactions influencing gene expression, with different regulatory outcomes that likely depend on several factors (e.g., regulatory elements within the interaction, the chromatin state, the bindings of transcription factors, and the mRNA stability, among others)^40,42^.

By integrating these molecular layers, we identified Selenoprotein W (*Selenow*) as a promising candidate gene for an aging signature in brain. *Selenow* is involved in neuroprotection against oxidative stress^56,57^, as well as memory formation^58^ and inhibition of tau protein aggregation^59,60^. Our analysis revealed significant changes in both *Selenow*’s promoter interactions and its gene expression with age. At the promoter interactome level, *Selenow* exhibited an age-associated decrease in interactions within a region downstream, with an accompanying increase in interactions (Chicdiff p-value of 8 interacting regions < 0.0000001) within a specific genomic region of the chromosome 7 (chr7:14,522,438-14,639,621) (Figure 2F, and Table S6). This implies that, with age, a loop is generated between the *Selenow* promoter and this specific region. Moreover, RNA-seq analysis revealed a significant upregulation of *Selenow* expression in older mice (p-value = 0.012; LFC = 0.646) (Figure 2G). Taken together, these results suggest that the age-related upregulation of *Selenow* expression might be, at least in part, mediated by chromatin reorganization and the formation of a chromatin loop.

To broaden the context of our findings, we compared them with existing data on aging in mice. We limited our comparison to the only previous work performing Promoter Capture Hi-C in aging mice, which analyzed pre-B cells^61^. Our analysis revealed minimal overlap, with only 10 interactions showing significant changes with age in both neurons and pre-B cells (Figure S4A). Intriguingly, 7 out of these 10 shared interactions clustered within a specific region on chromosome 19, involving the *Kif11* promoter. In both neurons and pre-B cells, this region displayed increased contacts with aging (Figure S4B, and Table S7). Additionally, *Kif11* expression in the hippocampus exhibited downregulation with age (Figure S4C). *Kif11* is a kinesin that plays a role in formation of the mitotic spindle^62^, and has been linked with the maintenance of learning and memory in the Alzheimer’s disease^63^. These findings suggest that, despite cell type differences, the aging process might involve some common alterations in the chromatin interactome.

### Environmental Enrichment has an age-dependent impact on the chromatin interactome

To investigate whether external stimuli may influence and modulate chromatin interactions, we explored the impact of environmental enrichment (EE) on promoter interactions. First, we analyzed the impact of EE within each age group, distinguishing between EE in young mice (YEE vs young control) and EE in old mice (OEE vs old control) (Figure 3A). liCHi-C experiments were performed on samples from EE mice, and sequencing data was processed identically to the control samples. No significant differences were observed between the replicates for both control and EE samples using HiCUP and CHiCAGO statistics. This included the percentage of captured valid reads, the *cis*-*trans* proportions, the number of CHiCAGO-significant interactions, and the promoter-promoter interactions proportion (Figures 1C-F, S5A-D and Table S1).

**Figure 3.**
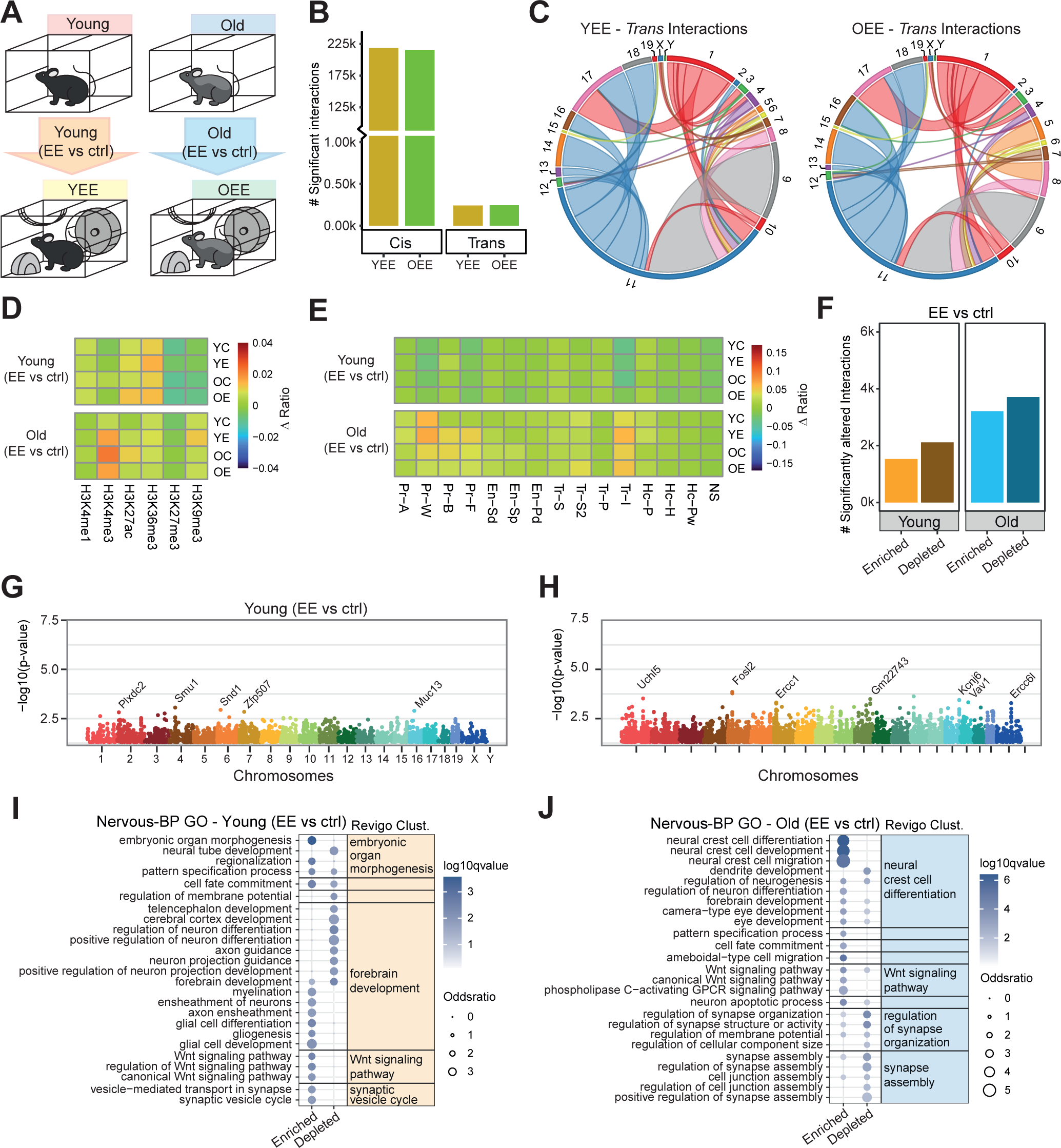
Promoter interactions alterations with the Environmental Enrichment in young mice and in old mice. **A.** Schematic of the environmental enrichment (EE) design. We distinguish between the exposure to environmental enrichment in young mice (EE in young), comparing young mice and young with environmental enrichment (YEE) mice; and the exposure to environmental enrichment in old mice (EE in old), comparing old mice and old with environmental enrichment mice (OEE). **B**. Bar plot illustrating the total number of CHiCAGO significant interactions in *trans* and in *cis* for YEE and OEE mice. 1000 (k). **C**. Chord plots representing the *trans* interaction proportion of each mice chromosome (outside number) for young mice (left) and old mice (right). **D, E**. Heatmaps representing histone mark (**D**) and chromatin states (**E**) enrichment analysis of significant interactions in *cis*. Color scale reflects the differential ratio (Δ ratio) with EE in young mice (top) and in old mice (bottom), and it was calculated subtracting the control conditions values from the EE conditions values. Six common histone posttranslational modifications, and the 15 chromatin states (see Methods section) defined by histone marks, from four mouse hippocampus datasets (the four experimental conditions: YC, YE, OC and OE) of a previous study^47^ were used in the enrichment analysis. **F**. Bar plot illustrating the total number of Chicdiff interactions in *cis* that change significantly (p-value < 0.05) with the EE in both young and old mice, distinguishing between enriched and depleted interactions. 1000 (k). **G, H**. Manhattan plot of differential interactions in *cis* (Chicdiff p-value < 0.05) with EE in young mice (**G**) and EE in old mice (**H**). **I, J**. Bubble plot represents Gene Ontology (GO) terms enrichment based on Nervous system-associated Biological Processes (Nervous-BP) for promoter genes with significantly enriched and depleted interactions (p-value < 0.05) with the EE in young mice (**I**) and in old mice (**J**). Bubble color intensity represents the statistical significance (-Log_10_ q-value) and dot size reflects odds ratio. Top significant GO terms were grouped following REVIGO clustering, which are named with the most representative term of the group.

We analyzed the number of significant interactions in *trans* and *cis* for YEE and OEE mice. For interactions in *trans*, we identified 241 CHiCAGO significant contacts in YEE mice and 245 in OEE mice (Figure 3B). Furthermore, YEE and OEE mice exhibited distinct patterns compared to their respective control conditions. For example, we observed a higher number of interactions between chromosomes 11 and 17 in YEE and an increase in interactions between chromosomes 5 and 8 in OEE (Figures 1C, 3C, S1G and S5E). The total number of significant *cis* interactions also remained comparable, with over 200,000 identified in both YEE (218,633) and OEE (215,871) mice (Figure 3B), with median linear distances of 263 kb and 250 kb respectively, with the distance distribution between OEE and old control conditions being significantly different (Wilcoxon p-value < 0.001) (Figure S5F). However, the number of *cis* contacts per promoter was significantly lower in EE conditions (YEE and OEE) than their corresponding control conditions (Figure S5G), which was also evident when exploring chromosomes individually (Figures 1E, S5H). Thus, exposure to EE revealed promoter interaction alterations in *cis*, but also in *trans* interactions.

Subsequently, we conducted histone and chromatin state enrichment analysis within *cis*-interacting regions for EE conditions, and we compared them with control conditions. While histone modifications exhibited minor variations in response to EE (Figure 3D), we did observe slight changes in specific chromatin states (Figure 3E). Interestingly, the enrichments for weak promoter and transcription initiation states exhibited opposing trends in young and old mice with EE. With EE in young mice, these states showed a slight decrease in enrichment, mirroring the trend observed with aging. Conversely, with EE in old mice, the enrichment for these states increased slightly (Figure 3E).

Differential analysis using Chicdiff revealed distinct patterns of interaction alterations upon EE (Figure 3F). With EE in young mice, we observed a higher number of significantly depleted interactions (2,116) compared to enriched interactions (1,531) (Figure 3F), including, for instance, *Smu1* and *Snd1* promoter interactions (Figure 3G). Similarly, with EE in old mice, we identified more depleted interactions (3,706) than enriched interactions (3,212) (Figure 3F). These altered interactions involved *Fosl2* and *Uchl5* promoters (Figure 3H). Notably, there were fewer significantly altered interactions observed under EE conditions compared to what we observed with aging (almost 10,000 changes; Figure 1H). This suggests that aging had a stronger impact on chromatin interactome than environmental enrichment, at least on hippocampal neurons.

Analyzing the possible relationship between the interaction changes upon EE and chromatin common histone marks, significantly altered interactions with EE were associated with changes on active histone marks in promoters, but with a subtle fold change magnitude. Interestingly, with EE in young, active histone marks followed the same trend observed with aging, while with EE in old, active histone marks had an opposite trend to the finding in aging (Figures S6A-F). Moreover, in the interacting regions, similar but slighter tendencies are observed (Figures S7A-F).

Gene ontology (GO) analysis of gene promoters with altered interactions upon EE revealed several genes that were associated with different biological processes in young and old mice. Some processes, such as Wnt signaling, were mainly associated with enriched interactions with EE in both young and old mice. However, other processes seemed to be associated with depleted interactions with EE in young mice, but with enriched interactions with EE in old mice, e.g. processes related to nervous system development and differentiation (Figures 3I, J, S5I, S5J and Table S2). Taken together, all these findings suggest that EE has a differential effect on chromatin contacts in young and old mice.

### The age-dependent effect of EE on chromatin interactions is recapitulated at the transcriptional level

Transcriptome analysis of the entire hippocampus, comparing control and EE samples, revealed a clear dominance of age as a factor influencing gene expression variation. Replicates clustered primarily by age, indicating that age-related differences overshadowed the effects of EE (Figure 4A). Differential expression analysis identified a higher number of genes significantly upregulated (455) or downregulated (444) with EE in young mice, compared to the old mice (177 upregulated, 167 downregulated). Remarkably, most differentially expressed genes (DEGs) were specific to either EE in young or in old mice, with the highest overlap (15 DEGs) occurring between upregulated genes in YEE mice and downregulated genes in OEE mice (Figures 4B, 4C). Moreover, GSEA analysis revealed an age-dependent effect of EE on specific pathways. For example, several pathways associated with extracellular matrix and collagen biosynthesis were upregulated with EE in young mice but downregulated with EE in old mice (Figure 4D and Table S4). Taken together, these findings strongly suggest that, similarly to the chromatin interactome, the impact of EE on gene expression is significantly influenced by the age of the organism, with more DEGs in young than old mice. Moreover, certain molecular pathways are differentially modulated by EE in young and old individuals.

**Figure 4.**
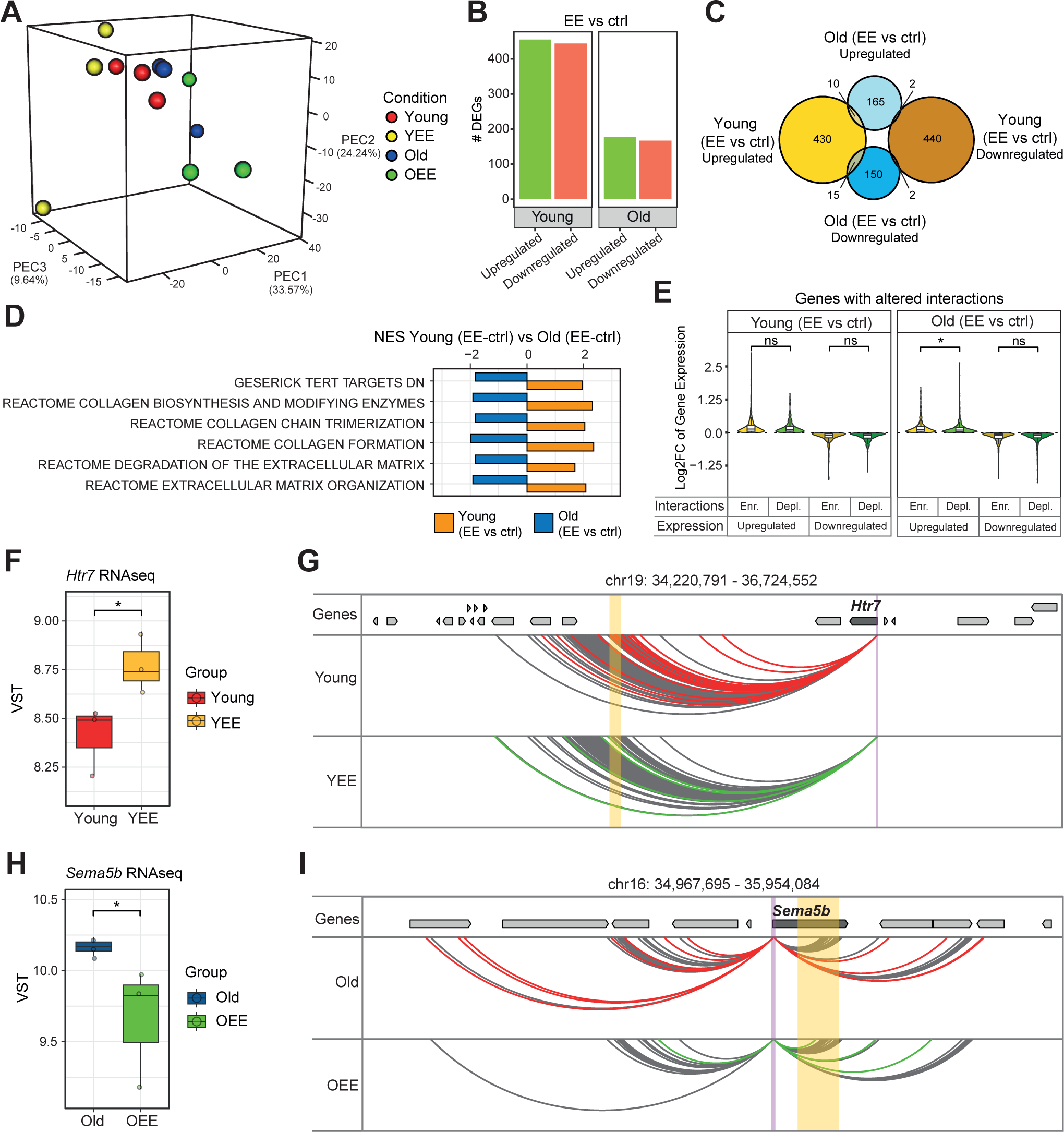
Environmental Enrichment effects at RNA level and integration with liCHi-C data. **A.** Principal Component Analysis (PCA) for all expressed genes of RNA-seq replicates in young, old, YEE and OEE conditions. The proportion of variance explained for the PC1, PC2 and PC3 components is indicated. **B**. Bar plot illustrating the number of Differentially Expressed Genes (DEGs) (p-value < 0.05) with EE in young mice (left) and EE in old mice (right), distinguishing between upregulated and downregulated genes. **C**. Venn diagram depicting the number of up- and down-regulated genes for EE in young mice and EE in old mice. **D**. Barplots reflecting the Normalized Enrichment Score (NES) of GSEA gene sets from M2 collection (MSigDB). Gene sets NES for EE in young mice and EE in old mice are shown (FDR q-value < 0.2). **E.** Violin plots showing the Log_2_ Fold Change of genes with altered interactions in response to EE in young mice (left) and in old mice (right). Upregulated and downregulated genes with EE are represented separately, comparing those with enriched or depleted interactions with EE. Significance calculated with Wilcoxon rank-sum test. **F**. Boxplots indicating the RNA-seq expression measurements (VST normalized data) for the *Htr7* gene across young and YEE mice. Dots denote individual mouse. **G**. *Htr7* promoter-centered interactions according to liCHi-C data in young and YEE conditions. Arcs represent CHiCAGO significant contacts, red arcs highlight those contacts in young mice that were lost in YEE mice, while green arcs highlight those contacts that are found in YEE mice but not in young mice. Purple shade depicts *Htr7* gene promoter and yellow shade depicts a region with significantly altered interactions. **H**. Boxplots indicating the RNA-seq expression measurements for the *Sema5b* gene across old and OEE mice. Dots denote individual mouse. **I**. *Sema5b* promoter-centered interactions according to liCHi-C data in old and OEE conditions. Arcs represent CHiCAGO significant contacts, red arcs highlight those contacts in old mice that were lost in OEE mice, while green arcs highlight those contacts that are found in OEE mice but not in old mice. Purple shade depicts *Sema5b* gene promoter and yellow shade depicts a region with significantly altered interactions. Arrows symbolize gene placement and orientation along the genomic window. ^ns^p-value > 0.05, *p-value < 0.05.

The integration of chromatin interactions and gene expression revealed that, in general terms, there is not a clear nexus between interaction and expression changes (Figures 4E, S3B, S3C). Also, most of the genes with significant changes in both interaction patterns and expression levels were specific to each age group (Figures S3B, S3C and Table S5). One example of them with EE in young mice was *Htr7*, a serotonin receptor belonging to the GPCR (G protein-coupled receptors) family that has been associated with cognitive and synaptic plasticity^64^. *Htr7* showed a significant increase in expression with EE in young mice (p-value = 0.016; LFC = 0.483) (Figure 4F), accompanied by a significant decrease in promoter interactions (Its most significantly altered interaction: p-value = 0.012; LFC = -0.271) (Figure 4G and Table S6). Conversely, expression of *Sema5b*, a gene involved in axon growth and reduction of synapse connections^65^, was downregulated with EE in old mice (p-value = 0.025; LFC = -0.534) (Figure 4H), accompanied by a significant decrease of its promoter interactions (Its most significantly altered interaction: p-value = 0.020; LFC = -0.175) (Figure 4I and Table S6). These findings highlight the age-dependent influence of EE on promoter interactions and gene expression of specific genes.

### Age-associated changes in the chromatin interactome are boosted by EE in young mice and partially reversed by EE in old mice

To explore the relationship between aging and EE, we compared these processes with one another (aging vs EE in young; aging vs EE in old). Around 4.2% of the interactions significantly altered with aging also exhibited significant changes with EE in young mice, while over 10% (10.817%) of these interactions altered with aging also changed significantly with EE in the old mice (Figure S8A). This supports the notion that EE might have a more substantial impact on the chromatin structure in older individuals compared to younger ones. Furthermore, the fold change of interactions revealed distinct patterns. In young mice, the direction of change (increase or decrease) in interactions upon EE generally mirrored the change tendency observed with aging (Figures 5A and S8B). However, in old mice, EE induced changes in interactions with the opposite trend compared to those observed with aging (Figures 5B and S8C).

**Figure 5.**
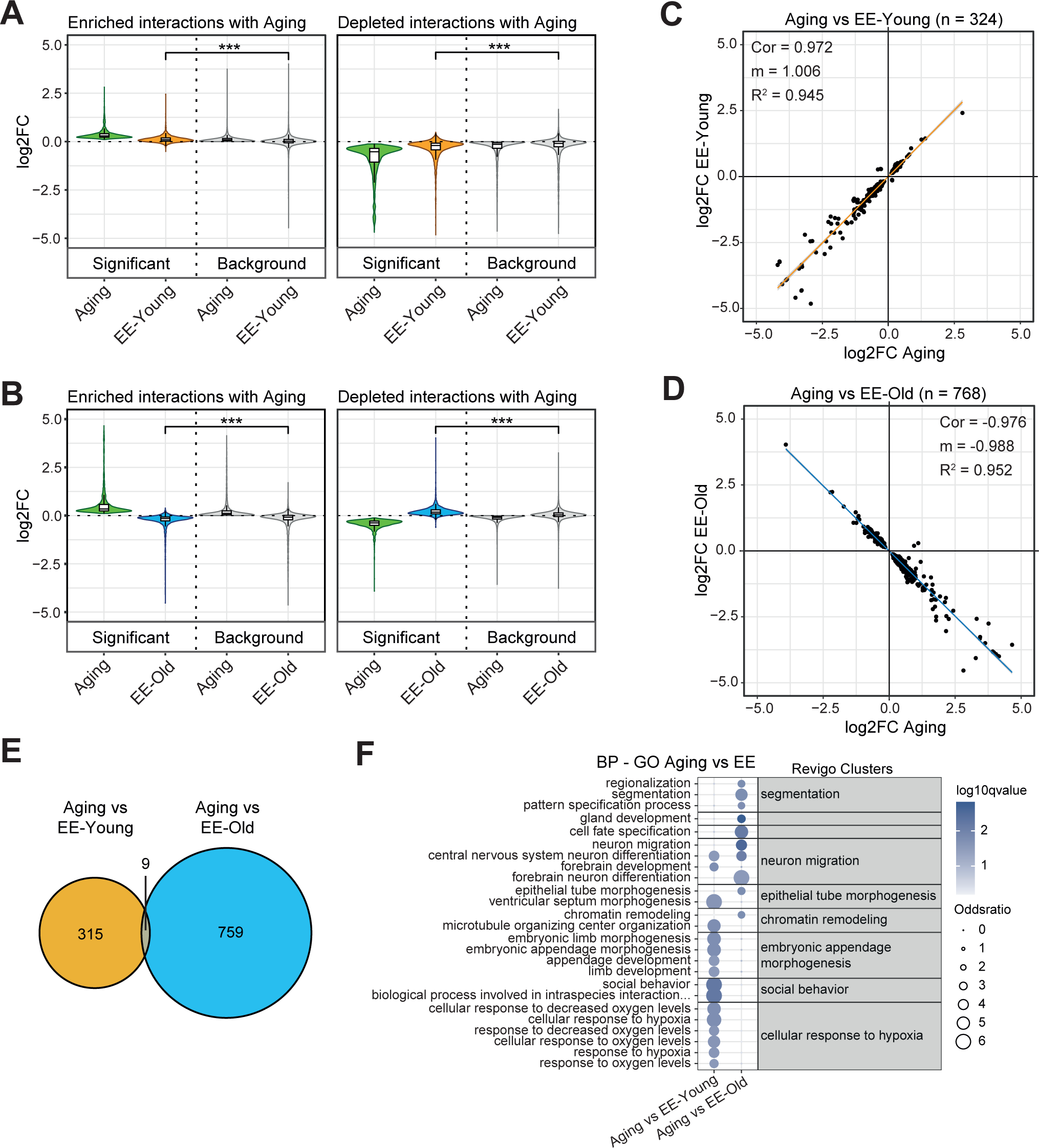
Comparative analysis between aging and Environmental Enrichment in both young and old mice at promoter chromatin interactome level. **A, B.** Violin plots showing the Log_2_ Fold Change for promoter interactions under two comparatives: with aging and EE in young mice (**A**), or with aging and EE in old mice (**B**). Enriched interactions (left) and depleted interactions (right) with aging were represented separately. Interactions significantly altered with aging (p-value < 0.05; Significant) were compared to those that were not significantly altered with aging (p- value > 0.05; Background). Significance calculated with Wilcoxon rank-sum test. ***p- value < 0.01. **C.** Scatter plot indicating a back-to-back comparison of Log_2_ Fold Change of significantly altered interactions for aging (p-value < 0.05) and for EE in young mice (p-value < 0.05) (Pearson’s correlation score = 0.972; p-value < 2.2e-16). Orange line indicates the regression line (slope and R-squared value are shown), and the gray shade represent the confidence interval. **D.** Scatter plot representing a back-to-back comparison of Log_2_ Fold Change of significantly altered interactions for aging (p-value < 0.05) and for EE in old mice (p-value < 0.05) (Pearson’s correlation score = -0.976; p- value < 2.2e-16). Blue line indicates regression line (slope and R-squared value are shown), and the gray shade represent the confidence interval. **E.** Venn diagram depicting the number of significant interactions (p-value < 0.05) for aging and EE in young mice (orange), or for aging and EE in old mice (blue). The number of overlapping interactions (significant for aging and both types of EE) is highlighted in grey. The overlap has a representation factor of 19.9 (p-value = 1.271 x 10^-9^; universe = 549588 interactions) using hypergeometric distribution. **F**. Bubble plot represents Gene Ontology (GO) terms enrichment based on all Biological Processes (BP) for gene promoter with altered interactions with aging (p-value < 0.05) and EE in young mice (p- value < 0.1), or with aging (p-value < 0.05) and EE in old mice (p-value < 0.1). Less restrictive significances for EE were used to be able to perform the GO analysis. Bubble color intensity represents the statistical significance (-Log_10_ q-value) and dot size reflects odds ratio. Top significant GO terms were grouped following REVIGO clustering, which are named with the most representative term of the group.

We performed further analysis focused on specific interactions that changed significantly with both aging and EE (Figures 5C, 5D). In young mice, these overlapping changes (n = 324) showed a positive correlation (Slope = 1.006; R² = 0.945) (Figure 5C), indicating that EE in young mice partially mimics the age-related trends in chromatin interactions. However, in old mice, the overlapping changes (n= 768) exhibited a negative correlation (slope = -0.988; R² = 0.952) (Figure 5D), demonstrating that EE tends to partially reverse the age-related changes in promoter interactions in old mice.

Only a small number of interactions (9 out of 1,083) displayed significant changes in all three processes: aging, EE in young and EE in old (Figure 5E). Furthermore, the genes with specific interaction changes for young mice or for old mice, were involved in distinct biological processes (Figure 5F). This indicates that those altered interactions are highly age-specific. In young mice, those gene promoters with altered interactions upon aging and EE were mainly associated with processes like social behaviour, response to hypoxia, and morphogenesis; while those altered in old mice were linked to processes such as segmentation, cell fate specification, or neuron migration and differentiation (Figure 5F and Table S2). Taken together, our findings highlight the differential effect of EE depending on age, with EE promoting age-related changes in promoter interactions in young mice, and partially counteracting aging-related alterations in old mice.

### The age-related differential effect of EE on the chromatin interactome is further mirrored at the level of gene expression

To investigate the relationship between aging and EE at transcriptional level, we conducted the same comparative analysis (aging vs EE in young; aging vs EE in old) with RNA-seq data. Gene set enrichment analysis (GSEA) identified several gene sets exhibiting opposing modulation with aging and EE in young mice (e.g., dopaminergic neurogenesis or telencephalon microglia), while others changed in the same direction (e.g., *FoxP3* targets) (Figure 6A and Table S4). In old mice, some gene sets exhibited similar variations (e.g., ribosomes and mitochondrial translation pathways), while several gene sets, most of them associated with GPCR signaling, showed differential expression shifts with aging and EE (Figure 6B and Table S4), which was also corroborated using GO terms (Figure S9A and Table S4). In fact, GPCRs play a crucial role in the synaptic organization and neurotransmission function^66^. Additional analysis using the LEIN markers dataset^67^, revealed enrichment of choroid plexus markers with EE in young mice but depletion with EE in old mice. Moreover, other brain markers (e.g., midbrain markers) that were depleted with aging, were enriched with EE in both young and old mice (Figure S9B and Table S4).

**Figure 6.**
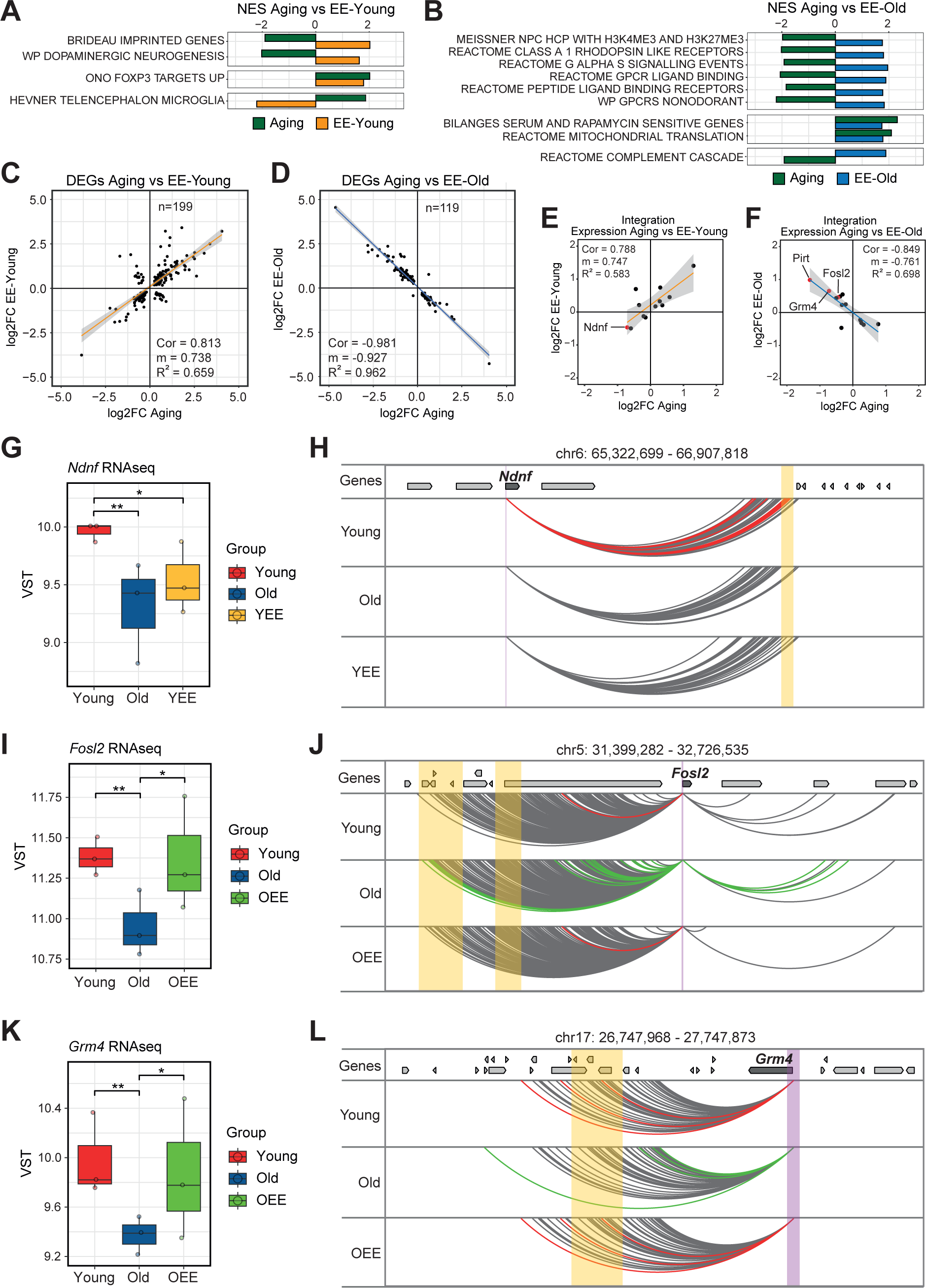
Gene expression differences between aging and the Environmental Enrichment in both young and old mice, and the integration with promoter interactions. **A, B**. Barplots reflecting the Normalized Enrichment Score (NES) of GSEA gene sets from M2 collection (MSigDB). Gene sets NES for aging (old vs young) and EE in young mice (**A**), or for aging and EE in old mice (**B**) are shown (FDR q-value < 0.05 for aging; FDR q-value < 0.2 for EE). Gene sets were grouped in clusters based on NES signs. **C**. Scatter plot indicating a back-to-back comparison of Log_2_ Fold Change of DEGs for aging (p-value < 0.05) and EE in young mice (p-value < 0.05) (Pearson’s correlation score = 0.813; p-value < 2.2e-16). Orange line indicates the regression line (slope and R-squared value are shown), and the gray shade represent the confidence interval. **D**. Scatter plot indicating a back-to-back comparison of Log_2_ Fold Change of DEGs for aging (p-value < 0.05) and EE in old mice (p-value < 0.05) (Pearson’s correlation score = -0.981; p-value < 2.2e-16). Blue line indicates the regression line (slope and R- squared value are shown), and the gray shade represent the confidence interval. **E**, **F**. Scatter plot indicating a back-to-back comparison of Log_2_ Fold Change of gene expression (p-value < 0.2) for aging and for EE in young mice, whose promoters also have significantly altered interactions (p-value < 0.05) with aging and EE in young mice (Pearson’s correlation score = 0.788 and p-value = 0.002) (**E**); or of gene expression (p-value < 0.2) for aging and EE in old mice, whose promoters also have significantly altered interactions (p-value < 0.05) with aging and EE in old mice (Pearson’s correlation score = -0.849 and p-value = 0.0001) (**F**). Color lines indicate regression lines (slopes and R-squared values are shown), and the gray shades represent the confidence intervals. Red dots highlight genes with a p-value < 0.05 for gene expression and interactions with both aging and EE in young mice. **G**. Boxplots indicating the RNA-seq expression measurements (VST normalized data) for the *Ndnf* gene across young, old and YEE mice. Dots denote individual mouse. **H**. *Ndnf* promoter-centered interactions according to liCHi-C data in young, old, and young with environmental enrichment (YEE) conditions. Arcs represent CHiCAGO significant contacts, red arcs highlight those contacts in young mice that were lost in old and YEE mice. Purple shade depicts the gene promoter and yellow shade depicts a region with significantly altered interactions between represented conditions (old vs young; YEE vs young). **I**. Boxplots indicating the RNA-seq expression measurements for *Fosl2* gene across young, old and OEE mice. Dots denote individual mouse. **J**. *Fosl2* promoter- centered interactions, according to liCHi-C data in young, old, and old with environmental enrichment (OEE) conditions. Arcs represent CHiCAGO significant contacts, red arcs highlight those contacts in young mice that were lost in old mice and gained in OEE mice, while green arcs highlight those contacts that are found only in old mice but not in young or OEE mice. Purple shade depicts the gene promoter and yellow shades depicts a region with significantly altered interactions between represented conditions (old vs young; OEE vs old). **K** and **L.** As in (**I** and **J**), respectively, but for *Grm4* gene and promoter. Arrows symbolize gene placement and orientation along the genomic window. ^ns^p-value < 0.05, *p-value < 0.05, **p-value < 0.01.

Further analysis that focused on DEGs for aging and EE, revealed that gene expression fold changes, as well as EE-specificity, followed the same trends we observed with chromatin interactions, i.e. in young mice, the direction of change (increase or decrease) in expression upon EE generally followed the tendency observed with aging, while in old mice, EE induced changes with the opposite trend observed with aging (Figures 6C, 6D and S9C-E). Remarkably, these tendencies were further corroborated by the integrative analysis of promoter interactions and gene expression, obtaining some genes with significantly altered interactions with aging and EE (p-value < 0.05), whose gene expression (p-value < 0.2) follow the trends described above (Figures 6E, 6F and Table S8). Notably, *Ndnf* was the only gene that met the condition of a 0.05 p-value threshold for both interaction and expression level for aging and EE in young mice (Figure 6E), while only *Fosl2*, *Grm4, and Pirt* met this criterion for aging and EE in old mice (Figure 6F).

The Neuron derived neurotrophic factor (*Ndnf*) is mainly expressed in neurons, and promotes neuronal growth, migration, and neurite outgrowth, among others^68^. At gene expression level, *Ndnf* was downregulated with both aging (p-value = 0.003; LFC = - 0.706) and EE in young mice (p-value = 0.021; LFC = -0.466) (Figure 6G). Nonetheless, *Ndnf* promoter interactions decreased with both aging and EE in young mice. Specifically, the most significantly altered interaction for *Ndnf* was reduced with both aging (p-value = 0.005; LFC = -0.532) and with EE in young mice (p-value = 0.015; LFC = -0.519) (Figure 6H, and Table S6).

In the comparison of aging and EE in old mice, *Fosl2* emerges as one of the examples for the partial reversal observed at interaction and expression level. *Fosl2* encodes a transcription factor involved in several processes such as cellular proliferation, differentiation and apoptosis ^69^. *Fosl2* expression was downregulated with aging (p-value = 0.003; LFC = -0.445) but was reversed with EE in old mice (p-value = 0.036; LFC = 0.460) (Figure 6I). Together, some of the *Fosl2* promoter contacts increased with aging, but decreased with EE in old mice. Particularly, its most significantly altered interaction with aging (p-value = 0.002; LFC = 0.426) was reverted by EE in old mice (p-value = 0.002; LFC = -0.444) (Figure 6J and Table S6).

Another example of the reversion of age-related changes with EE in old mice was *Grm4,* a GPCR for the neurotransmitter L-glutamate^70^ that has been associated with learning^71^, and synapse remodelling in memory consolidation^72^. *Grm4* expression was downregulated during aging (p-value = 0.001; LFC = -0.723) and upregulated with EE in old mice (p-value = 0.028; LFC = 0.654) (Figure 6K). Some *Grm4* promoter interactions decreased with aging (its most significantly altered interaction: p-value = 0.006; LFC = -0.324), while they were reverted with EE in old mice (p-value = 0.003; LFC = 0.370) (Figure 6L and Table S6).

Taken together, our findings reveal an age-dependent effect of EE in hippocampal neurons. In young mice, EE seems to stimulate some age-related changes in promoter interactome and gene expression, possibly associated with an early maturation of the hippocampus. Conversely, in old mice, some of the interactome and transcriptome changes identified with aging can be reversed by EE, potentially indicating a partial rejuvenation of the hippocampal neurons.

## DISCUSSION

The aging process involves an accumulation of degenerative phenomena due to molecular alterations over time. However, further research is needed to unravel the complexities of the molecular mechanisms underlying aging, especially in the context of the 3D chromatin interactome. To address this issue, we used a mouse model comprising both young and old individuals and performed liCHi-C method in dorsal hippocampal neurons. To complement this data with functional insights, we conducted an RNA-seq analysis of bulk dorsal hippocampus. Therefore, we characterized age-associated changes at promoter interactions and gene expression, which appear to be mainly related with certain biological processes. For instance, with aging, we describe a chromatin loop formation at *Selenow* promoter, which possibly involved in *Selenow* expression increase with aging, suggesting a neuroprotective role during aging against oxidative stress^56,57^ and Alzheimer’s disease^59,60^.

In this study, we also investigated whether the alterations observed in the chromatin interactome and transcriptome with age could be influenced by external stimuli. To explore the possible modulatory effect of an external factor, we exposed young and old mice to environmental enrichment (EE) and analyzed their hippocampi at the molecular level. Therefore, we determined altered interactions and DEGs with the EE, strengthening the notion that external stimuli can modulate chromatin interactions and gene expression in brain^30^. Our findings demonstrated that cognitive and physical stimulation have an age-dependent impact on brain at these molecular layers.

After describing the molecular landscape in the context of aging and environmental enrichment, we aimed to elucidate the relationship between both processes. Remarkably, we observed a strong negative correlation between those interactions that were significantly altered by aging and by EE in old mice, but a strong positive correlation between interactions altered by aging and by EE in young mice. Furthermore, a similar pattern was observed at gene expression level.

With EE in old individuals, we propose that we may be describing a partial rejuvenation of the hippocampus at the promoter interactome level associated with environmental enrichment, which results in the brain resembling a younger brain at molecular level, consistent with our and others’ previous findings at other molecular layers^46,47^. For instance, we observed this rejuvenation at specific gene promoters, such as the *Fosl2* and *Grm4* promoters. *Fosl2* alterations at Interactome and transcriptome levels, possibly linked to a higher risk for Parkinson’s disease^73^, could be partially reversed with EE, fostering a partial rejuvenation of the brain. Additionally, the G protein-coupled receptors (GPCR), which have been described to be modulated by environmental factors^30^, also seems to be impaired with aging and partially counteracted with EE in old mice. Especially, the *Grm4* downregulation with aging^74–76^ and its associated promoter interaction alterations, can be partially reversed by an environmental enrichment, leading to neuronal rejuvenation.

Regarding the induction of aging-like alterations upon EE in young mice, we can speculate that some of the molecular alterations that occur in the hippocampus during aging may also be induced by cognitive and physical stimulation, promoting an early maturation of the brain in young individuals. Brain maturation is produced by the accumulation of experiences, learning, and memory consolidation. Therefore, external stimuli that fosters the acquisition of these experiences could promote this process^77^. Furthermore, we suggest that an early maturation of the hippocampus may lead to an accelerated reduction of cell proliferation and differentiation, as well as a decline of their associated markers, such as *Ndnf,* similarly to other neurotrophic factors (e.g. *Bdnf*)^78^. However, further studies are needed to elucidate the effect of environmental enrichment on the early-life brain.

We described different patterns and trends of promoter interactions and gene expression associated with a partial rejuvenation and maturation of the hippocampus. This was achieved using different starting material for the analysis of chromatin interactome of NeuN^+^ hippocampal neurons and transcriptome of whole hippocampus (where over 60% of cells are NeuN^+^ neurons), due to technical constraints. Future research using single-cell approaches may facilitate the interpretation of the contribution of cell heterogeneity in the context of the different molecular layers, and obtaining a comprehensive characterization of partial rejuvenation and maturation of the hippocampus.

In summary, we have characterized alterations in the chromatin interactome upon aging in hippocampal neurons, focusing on promoter interactions, and their functional implications at the transcription level. Moreover, we reveal an age-dependent effect of an external factor (Environmental Enrichment) that leads to partial neuronal rejuvenation in old mice and to potentially early maturation of hippocampus in young individuals. Further research will shed light about the effects of different external stimuli on young brains and elucidate their rejuvenating potential for aged hippocampi.

## METHODS

### Mouse Model

A total of 48 male C57Bl/6J mice (*Mus musculus*), 9 weeks old (24 young mice) and 17 months old (24 old mice), were supplied by Janvier Labs. Mice from each age group were randomly assigned to different experimental groups: 12 to control conditions and 12 exposed to Environmental Enrichment for 2 months. Upon arrival at the Animal Facilities of the University of Oviedo, 5 or 6 animals were housed per cage, and they were acclimated for 2 weeks. Mice were maintained under standard conditions, including 12-hours light/dark cycles, controlled temperature (20-22°C), and *ad libitum* access to food and water, in accordance with the European Union Directive 2010/63/EU. The animals were euthanized by cervical dislocation, and the dorsal hippocampi were isolated from the mouse brains. All procedures were performed in accordance with the Spanish Guidelines (Real Decreto 53/2013) based on the European Union Directive 2010/63/EU for animal research, and were approved by The Research Ethics Committee of the University of Oviedo (PROAE 08/2021).

### Environmental Enrichment

Young and old mice exposed to environmental enrichment were housed in large PVC cages (Eurostandard Type IV, Biosis, Catalog #1354G, 598x380x250 mm) with tunnels, swings, wheels, ramps, houses, and different toys and objects of different shapes, sizes, and textures, randomly distributed throughout the cage. These objects were exchanged between cages every 2-3 days to modify the environmental stimuli and expose the animals to different mice odors. Also, bedding materials were changed every 1-2 weeks. Animals of control group were housed in the same room in standard-size cages (Eurostandard Type II L, Biosis, Catalog #1284L, 365x207x140 mm) without access to the objects and toys.

### Nuclei extraction and fixation

Hippocampi were homogenised with a syringe in cold NEB (Nuclei Extraction Buffer: 0.32 M sucrose, 25 mM KCl, 5 mM MgCl2, 20 mM Tris-HCl pH 7.5) with Triton X-100 (Sigma-Aldrich, Catalog #T8787) to a final concentration of 0.5%. The homogenates were then incubated for 15 minutes at 4°C followed by centrifugation at 3,200 xg for 5 minutes at 4°C, and the supernatants were discarded.

Samples were resuspended in 1x PBS containing 2% formaldehyde (Agar Scientifics, Catalog #R1026) and incubated in a rocker for 10 min at room temperature. Formaldehyde was quenched with glycline to a final concentration of 0.125 M and incubated for 5 min at room temperature followed by 15 min at 4°C. Subsequently, the samples were washed with cold 1x PBS.

### Neuronal Nuclei Purification

Neuronal nuclei were purified using fluorescence-activated nuclei sorting (FANS). Briefly, samples were stained with 7-AAD staining solution (Biolegend, Catalog #420404; diluted 1:40, 5 µl/test) and Alexa Fluor®488-conjugated Anti-NeuN Antibody (clone A60, Merck-Millipore, Catalog #MAB377X; diluted 1:2000), and then incubated overnight at 4°C. Stained samples were filtered through a 35 µm nylon mesh (Falcon, Catalog #352235), and FANS was performed using MoFlo XDP High-Speed Cell Sorter (Beckman Coulter, Catalog #ML99030) with Summit software (v5.2.0.7477). Neuronal nuclei were sorted intro DNA LoBind 1.5-ml tubes (Eppendorf, Catalog #022431021), flash-frozen in dry-ice and stored at -80 °C. For more details about sorting gates, see Figure S1.

### Low input Promoter Capture Hi-C library preparation and sequencing

Neuronal nuclei samples from 9 mice per experimental group were used in low input Promoter Capture Hi-C (liCHi-C)^79^. Nuclei were thawed and centrifugated (1000xg at 4°C for 10 min), and the supernatant was discarded. Then pelleted nuclei were resuspended in ice-cold lysis buffer (10 mM Tris-HCl pH 8.0, 10 mM NaCl, 0.2% IGEPAL CA630 and 1× cOmplete EDTA-free protease inhibitor cocktail). Samples from 3 mice were pooled to obtain 10^6^ nuclei per replicate, and liCHi-C was carried out with 3 replicates per experimental group.

Briefly, chromatin was digested overnight with HindIII enzyme (100 U/μl, New England Biolabs, Ref. #R0104T). Then, the cohesive ends of the restriction fragments were filled-in with dGTP, dTTP, dCTT and biotin-14-dATP (Invitrogen, Catalog #19524-016) using klenow polymerase (5 U/μl, New England Biolabs, Catalog #M0210L). Blunt ends of the fragments were ligated with T4 DNA Ligase (1U/µl, Invitrogen, Catalog #15224- 025), and chromatin was decrosslinked overnight with proteinase K at 10 mg/ml (Roche, Catalog #03115879001). DNA was isolated using phenol:chloroform:isoamyl alcohol 25:24:1 (Sigma, Catalog #P3803) extraction, and quantified using Qubit 3.0 fluorometer (Invitrogen).

DNA was sheared to an average size of 400 bp using Covaris M220 focused-ultrasonicator. Biotinylated ligated products were pulled down using Dynabeads MyOne streptavidin C1 paramagnetic beads (Thermo Fisher, Catalog #65001). DNA was repaired and fragment ends were dATP-tailed and ligated to PE Illumina adapters (Table S9). The library was then amplified for 8-9 cycles by PCR, and purified by double-sided selection using SPRI beads (0.4-1 volumes, CleanNA, Catalog #CNGS- 0050). DNA concentration was determined using an Agilent 2200 Tapestation with D1000 ScreenTape Analysis.

Ligation products containing promoters were captured using the SureSelectXT Target Enrichment System for the Illumina Platform (Agilent Technologies), following manufacturer’s instructions. The captured library was amplified for 4 cycles by PCR and purified using SPRI beads (0.9 volumes). Neuronal liCHi-C libraries were paired-end sequenced by BGI Genomics using DNBseq 100 + 100PE platform.

### LiCHi-C processing

Paired-end reads were processed using HiCUP^50^ (v0.8.3). First, the reference genome (*Mus musculus* GRCm38/mm10, Ensembl release 100) was indexed by Bowtie2^80^ (v2.4.1) and computationally digested using the HiCUP Digester tool with HindIII restrictions sequences. Subsequently, reads were truncated using the HiCUP Truncater tool and aligned individually to the digested reference genome using the HiCUP Mapper tool. Experimental artifacts and duplicated reads were removed using the HiCUP Filter and Deduplicator tools. Paired reads that did not overlap with captured fragments in either case, were filtered out. Consequently, we retained only unique valid captured reads, allowing for the calculation of capture efficiency. Library statistics for all samples are presented in Table S1.

### LiCHi-C interaction calling

Promoter interactions were called using CHiCAGO R package^51,81^ (v1.18.0) with default parameters. CHiCAGO implements a statistical model with two components: biological noise (based on detected random interactions) and technical noise (based on experimental artifacts); along with normalization and multiple testing methods. Samples were analyzed individually, as well as merged samples, to increase sensitivity between experimental conditions. Unlocalised and unplaced scaffolds were filtered. Interactions are considered significant if they have a CHiCAGO score ≥ 5. Interaction statistics for all samples are presented in Table S1.

### Histone and Chromatin state enrichment

Chip-seq datasets for hippocampus tissue from a previous study^47^ with four experimental conditions (YC, YE, OC, OE) were used for enrichment analyses of histone marks and chromatin states in the interacting regions. We used the 15 chromatin states established in that work: Promoter-Active (Pr-A), Promoter Weak (Pr- W), Promoter Bivalent (Pr-B), Promoter Flanking Region (Pr-F), Enhancer Strong TSS-distal (En-Sd), Enhancer Strong TSS-proximal (En-Sp), Enhancer Poised TSS-distal (En-Pd), Transcription Strong (Tr-S), Transcription Strong-2 (Tr-S2), Transcription Permissive (Tr-P), Transcription Initiation (Tr-I), Heterochromatin Polycomb-associated (Hc-P), Heterochromatin H3K9me3-associated (Hc-H), Heterochromatin Polycomb-associated weak (Hc-Pw) and No significant signal (Ns). Enrichments were carried out using CHiCAGO significant interactions in *cis*, and the peakEnrichment4Features function of the CHiCAGO R package. Ratios were calculated by comparing significant interaction overlaps and random sampling overlaps, and differential ratios (Δ Ratio) were calculated by subtracting control condition from the other condition. Finally, enrichments were visualized with the pheatmap package (v1.0.12).

### Differential analysis of interactions

Differential analysis between experimental conditions were computed using Chicdiff R package^53^ (v0.6) with the default parameters. Chicdiff utilizes CHiCAGO data converted by makePeakMatrix CHiCAGO tool that spreads the “Other End” region generating an interaction peak matrix. The analysis was carried out on interactions with a CHiCAGO score cutoff greater than 5 on merged samples data for either of the compared conditions. We obtain Fold Change and p-value information of differential interactions between conditions. Interaction landscape with CHiCAGO and Chicdiff tracks was plotted with plotdiffBaits function.

### Differential analysis of histone marks

Chip-seq datasets for hippocampus tissue from a previous study^47^ with four experimental conditions (YC, YE, OC, OE) were used in the differential analysis of common histone marks (H3K27ac, H3K4me3, H3K4me1, H3K36me3, H3K27me3, H3K9me3). Normalization and differential analysis were performed using R/Bioconductor package DiffBind^82^ (v3.0.15). Then, significant histone marks peaks (p-value < 0.05) were annotated to promoters and interacting regions displaying significantly enriched or depleted interactions, using the R/Bioconductor package GenomicRanges^83^ (v1.54.1).

### Gene ontologies enrichment analysis

We used the clusterProfiler^84^ (v4.10.0) R package to determine the enrichment of Gene Ontology (GO) term for biological processes (BP) with gene promoters exhibiting significantly altered interactions. Specific parameters were: org.Mm.eg.db for the organism and Benjamini-Hochberg for the method for adjustment. Ontology terms with less than 15 genes or more than 2000 genes were filtered out. A q-value threshold of 0.05 was applied, and results with odds ratio greater than 6 were limited to 6. GO term enrichment was performed using all available BP-GO terms or focusing on Nervous system-associated BP-GO terms. The BP-GO terms most closely associated with the nervous system were selected and updated from Neural-Immune Gene Ontology (NIGO)^85^ and Synaptic Gene Ontologies (SynGO)^86^ subsets (Table S10). For captured fragments associated with multiple gene promoters, we prioritize protein-coding genes for GO analysis. All the genes in the capture design were considered the universe, and odds ratio was calculated comparing a given gene dataset to the universe. For clearer visualization, similar GO terms were grouped using an in-house pipeline based on the REVIGO clustering algorithm from the rrvgo^87^ (v1.14.1) R package.

### RNA Extraction, library preparation and sequencing

Total RNA from fresh-frozen dorsal hippocampi was extracted from 3 mice per group (three biological replicates per condition) using RNeasy Mini Kit (Qiagen, Catalog #74104) following manufacturer’s protocol, and including DNase treatment. RNA concentration was quantified using Qubit RNA HS Assay kit (Thermo Fisher Scientific, Catalog #Q32852) and RNA integrity assessment was performed using Agilent 5300 Fragment Analyzer (Agilent Technologies). Library preparation was conducted with NEBNext Ultra II RNA Library Prep Kit (NEB, Catalog #E7770), including polyA selection, following manufacturer’s instructions. Finally, the mRNA-seq libraries were sequenced, generating 150 bp paired-end reads on an Illumina NovaSeq 6000 system.

### RNA-seq processing

The preprocessing and quality control of FASTQ files were conducted using fastp^88^ (v0.20.1) with the options: -r -M 10 -l 20 -p -x --adapter_fasta. Salmon^89^ (v1.5.0) was used for transcript-level quantification using the following options: --libType A -- validateMappings --seqBias --gcBias. For the pseudo-alignment, a decoy-aware index gentrome was constructed from the mm10 genome and transcriptome. All the downstream analysis and processing were carried out within R: transcript-level RNA-seq quantification files from Salmon were imported into R and aggregated to the gene level through the R/Bioconductor package tximport^90^ (v1.14.2). Ensembl IDs were annotated to gene symbols using the R/Bioconductor package biomaRt^91^ (v2.42.0). mapping details are presented in Table S3.

### Differential gene expression analysis

After aggregating transcript-level estimated counts to the gene level with tximport (v1.14.2), low-expression genes were filtered out with the filterByExpr() function of the R/Bioconductor package edgeR^92^ (v3.28.1). Subsequently, differentially expressed genes (DEGs) were identified from filtered gene-level estimated counts using the R/Bioconductor package DESeq2^93^ (v1.26.0) with generalized linear models. Differential analyses were performed for three comparisons: Aging (old vs young), EE in young mice (YEE vs young) and EE in old mice (OEE vs old). DEGs were determined with a p-value significance level of 0.05. Visualization of results as well as other analyses (e.g., PCA or GSEA analysis) were conducted using VST-normalized values, obtained with the vst function of DESeq2.

### Principal component analysis

Principal component analysis (PCA) was conducted using the plot3d function of rgl package (v1.2.1) with default parameters. PCA was performed with variance-stabilizing transformed (VST) expression of all genes detected (18,715).

### Gene set enrichment analysis

Gene Set Enrichment Analysis (GSEA)^55^ (v4.3.2) was performed using the variance-stabilizing transformed (VST) expression data, and the M2 (curated gene sets) and M5 (ontology gene sets) collections from the mouse MSigDB^94^ v2023.1 (Molecular signatures database). Significant gene sets were selected with an FDR significance level of 0.05 for aging, while a level of 0.2 was used for environmental enrichment (EE) in young and old mice, in order to perform the comparisons. LEIN markets dataset^67^ was used for a specific enrichment analysis of mouse brain markers.

### Data visualization

Result plots were generated using the ggplot2 (v3.4.2) R package, as well as the eulerr package (v7.0.2) for Venn diagram. An in-house pipeline was employed to create virtual 4C plots of individual gene promoters. To visualize significant interactions (CHiCAGO score ≥ 5) and genomic regions with significantly altered interactions (Chicdiff), as well as to generate chord plots and circle plots, the WashU Epigenome Browser^95^ (v54.0.3) was used.

## Data availability

All data are available in the main text or the supplementary materials. Additionally, the raw liCHi-C and RNA sequencing data are available in the European Nucleotide Archive (ENA), and can be accessed with the accession number: PRJEB75484. Chromatin state annotations from a previous study^47^ are available in a Zenodo repository, at https://doi.org/10.5281/zenodo.8372431. PCHi-C data for pre-B cells of young and old C57BL/6 male mice^61^, are publicly available in GEO repository, at GSE109671. We performed PCHi-C processing and downstream analysis using their raw sequencing data under the same conditions as with our liCHi-C data.

## ACKNOWLEDGEMENTS AND FUNDING

We are grateful to the Scientific and Technical Services (STSs) of the University of Oviedo, including the Animal Facilities of the University of Oviedo, and especially to Ana Cristina Salas Bustamante for her positive contribution to the project. This work was supported by the Spanish Association Against Cancer (PROYE18061FERN to MFF), the Asturias Government (PCTI) co-funding 2018-2023/FEDER (IDI/2018/146 and IDI/2021/000077 to MFF), the Spanish Institute of Health Carlos III (Plan Nacional de I+D+I) co-funding FEDER (PI21/01067 to MFF and AFF), the Spanish Association Against Cancer (PRYGN235109FERN to MFF), CIBERER Acciones Cooperativas y Complementarias Intramurales (ACCI20-34-U766 to MFF), the Spanish Institute of Health Carlos III (COV00624 to JRT and MFF), ISPA and the Asociación Galbán (2021- 052-INTRAMUR GALBAN-GOURR to RGU and 2023-165-GALBAN-TEVAJ to JRT), ISPA-Jannsen (2021-048-INTRAMURAL NOV-TEVAR to JRT), Spanish National Research Council (202020E092 to MFF), and the European Commission NextGenerationEU, through CSIC’s Global Health Platform (PTI Salud Global) and the Spanish Ministry of Science and Innovation through the Recovery, Transformation and Resilience Plan (SGL2021-03-39 and SGL2021-03-040). The laboratory of BMJ is supported by FEDER/Spanish Ministry of Science and Innovation (PID2021- 125277OB-I00). JGV is supported by the Spanish Ministry of Universities (FPU20/04659). RGU is supported by the Centro de Investigación Biomédica en Red (CIBER). JRT is supported by a Ramon y Cajal contract from the Spanish Ministry of Science and Innovation (RYC2021-031799-I). RFP (BP17-114) and PSO (BP17-165) are supported by the Severo Ochoa program. AR is supported by Spanish National Research Council (SOLAUT_00038505 SGL2103040). JMK is supported by Horizon Europe Framework Programme (HORIZON-HLTH-2022-STAYHLTH-01-two-stage, ref. 101080219). LTD is supported by the FPI Fellowship (PRE2019-088005). LR is supported by an AGAUR FI fellowship (2019FI-B00017). BMJ is supported by the Spanish Health Institute Carlos III (CP22/00127). We also acknowledge support from the Institute of Oncology of Asturias (IUOPA, supported by Obra Social Cajastur Liberbank, Spain), the Health Research Institute of Asturias (ISPA-FINBA) and Consorcio Centro de Investigación Biomédica en Red de Enfermedades Raras (CIBERER-ISCIII).

## AUTHOR CONTRIBUTIONS

J.G.V., R.G.U., A.F.F. and M.F.F. conceived, coordinated, and supervised the study. J.G.V. and R.G.U. designed all aspects of research, performed environmental enrichment experiments, processed samples, analyzed and interpreted the data, and wrote the manuscript. A.F.F., M.F.F., J.M.K. and J.L.T. participated in drafting the manuscript. A.P., P.S.O., A.R. and M.R.S. performed environmental enrichment experiments. J.R.T., R.F.P. and C.A.D. performed computational analyses and interpreted the data. L.T.D., L.R. and B.M.J. performed liCHi-C experiments. All authors revised, read, and approved the final manuscript.

## DECLARATION OF INTERESTS

The authors declare no competing interests.

## Notes

### Competing Interest Statement

The authors have declared no competing interest.

### Summary of Updates

Figures 1-6 have been adjusted to prevent overlap with the header in the PDF.

