## Supplemental Material for "Age-Dependent Maturation and Rejuvenation of the Neural 3D Chromatin Interactome in Enriched Environments"

### SUPPLEMENTARY FIGURES

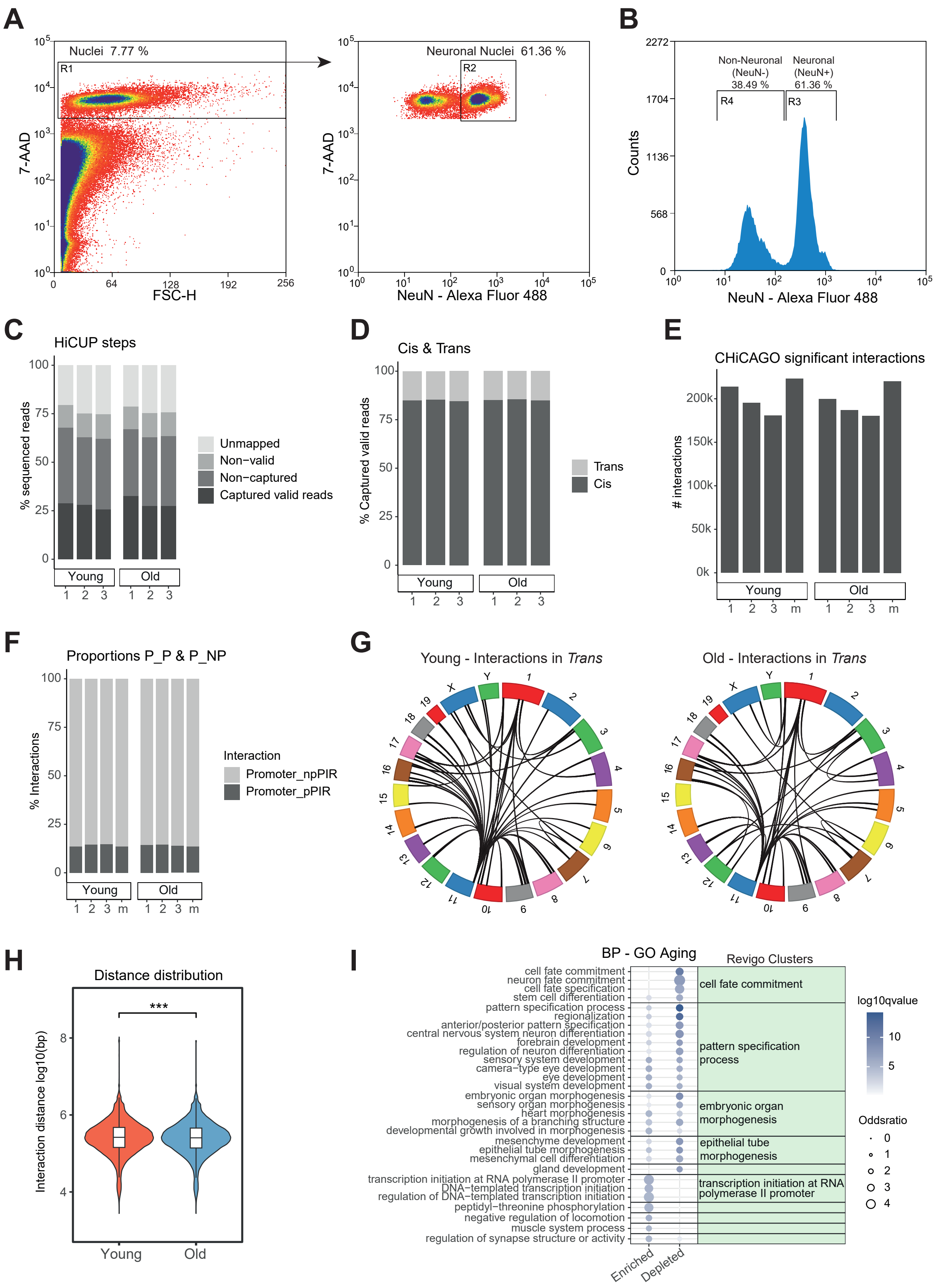

#### Figure S1. liCHi-C processing and analysis for young and old mice.

**A.** Schematic FANS sequential sorting gates to isolate neuronal nuclei. **B**. Histogram representing the proportion of $NeuN^{+}$ and $NeuN^{-}$ nuclei after FANS. **C**. Stacked barplots representing proportion of reads passed through the different steps of HiCUP in young and old liCHi-C replicates. **D**. Stacked barplots illustrating the cis-trans interactions proportion (in relative percentage) of captured valid reads in young and old liCHi-C replicates. **E**. Barplots showing the total number of CHiCAGO significant interactions (score > 5) of young and old liCHi-C replicates and merged samples (m). 1000 (k). **F**. Stacked barplots indicating the proportion of promoter-npPIR significant interactions (promoter and non-promoter PIR) and promoter-pPIR significant interactions (promoter and promoter PIR) of young and old liCHi-C replicates and merged samples (m). PIR: Promoter Interacting Region. **G**. Circle plots representing *trans* interaction between each mice chromosome for young mice (left) and old mice (right). **H**. Violin plots showing the distance distribution ($Log_{10}$ base pairs) of CHiCAGO significant interactions of merged samples for young and old conditions. Young and old distributions were compared using the Wilcoxon rank-sum test (p-value < 2.2e-16). ***p-value < 0.001. **I.** Bubble plot represents Gene Ontology (GO) terms enrichment based on all Biological Processes (BP) for promoter genes with significantly enriched and depleted interactions (p-value < 0.05) with aging (old vs young). Bubble color intensity represents the statistical significance (-$Log_{10}$ q-value) and dot size reflects odds ratio. Top significant GO terms were grouped following REVIGO clustering, which are named with the most representative term of the group.

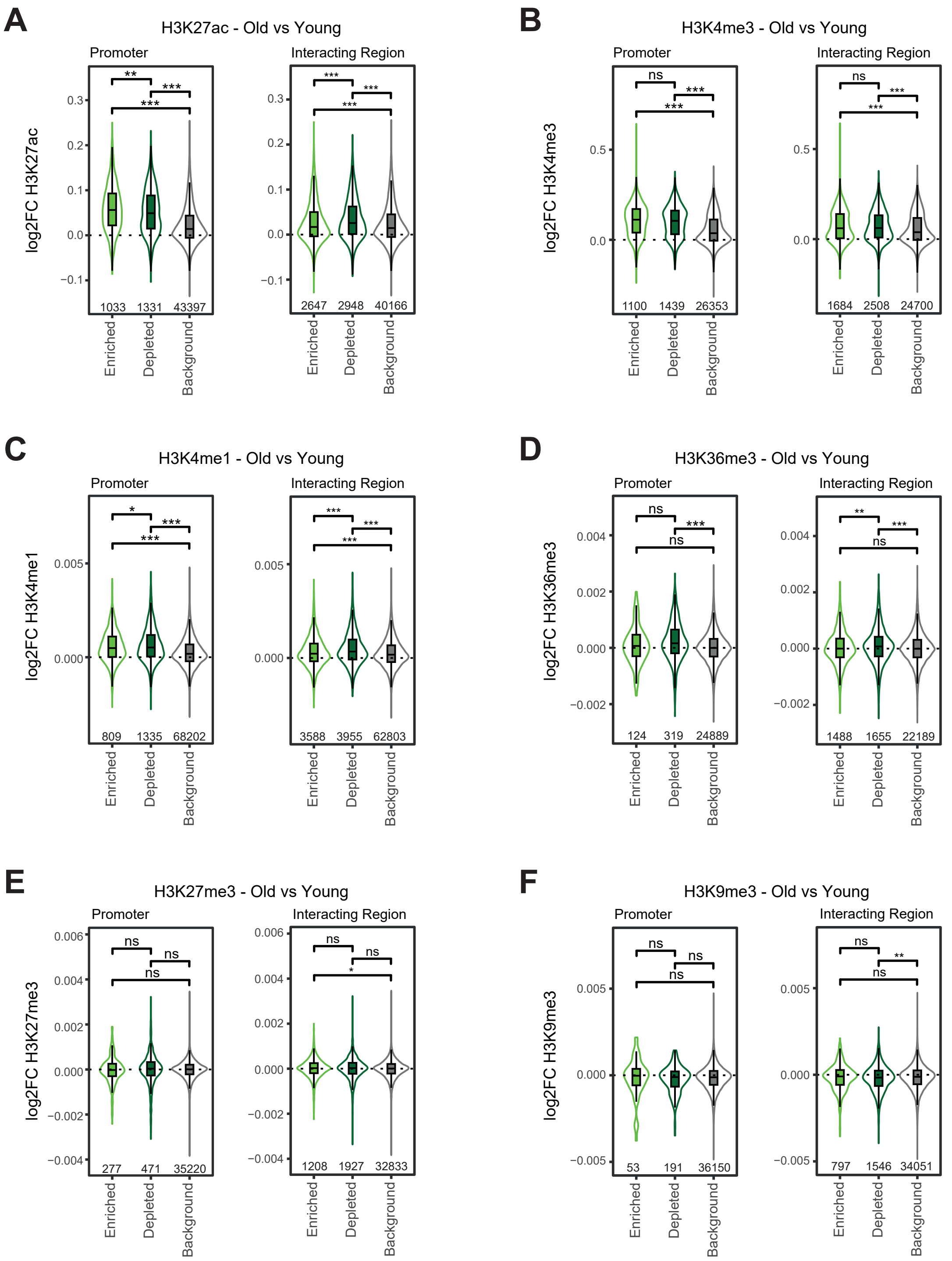


**Figure S2. Histone marks peaks of altered interactions with aging.**

Violin plots showing the differential ($Log_2$ Fold Change) with aging (old vs young) of histone marks from a previous study[47]: H3K27ac (**A**), H3K4me3 (**B**), H3K4me1 (**C**), H3K36me3 (**D**), H3K27me3 (**E**), H3K9me3 (**F**). Histone mark peaks were analyzed at promoter regions and interacting regions, differentiating between peaks associated with significantly enriched interactions (p-value < 0.05), with significantly depleted interactions (p-value < 0.05) and with non-significantly altered interactions (p-value > 0.05; background). Sample sizes are shown inside the plots. Wilcoxon rank-sum test was used to test whether the distribution histone marks $log_2FC$ differed significantly between groups. $^{ns}$p-value > 0.05, *p-value < 0.05, **p-value < 0.01, ***p-value < 0.001.

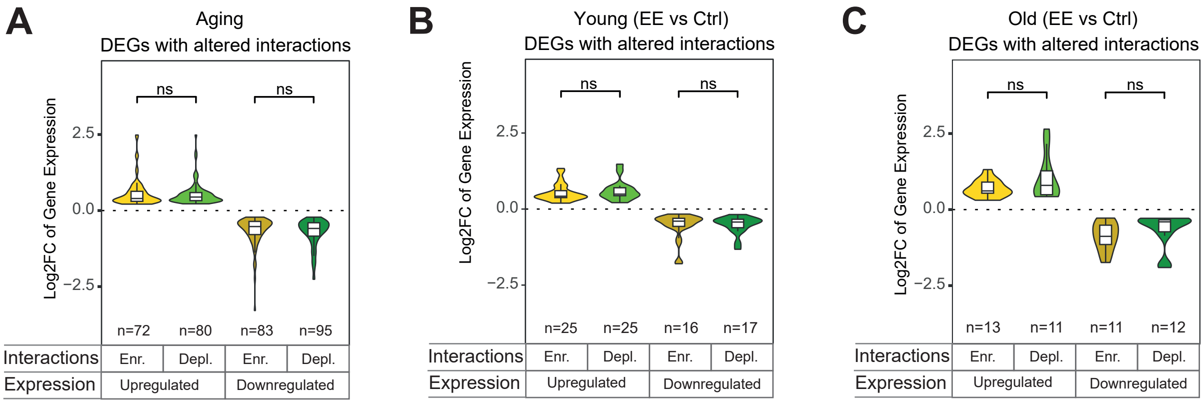


**Figure S3. liCHi-C and RNA-seq integration.**

Violin plots showing the $Log_2$ Fold Change of Differentially Expressed Genes (DEGs; p-value < 0.05) with altered interactions, with aging (**A**), EE in young mice (**B**) or EE in old mice (**C**). Upregulated and downregulated DEGs are represented separately, comparing those with enriched or depleted interactions. The number of DEGs for each combination is shown. Significance calculated with Wilcoxon rank-sum test. $^{ns}$p-value > 0.05.

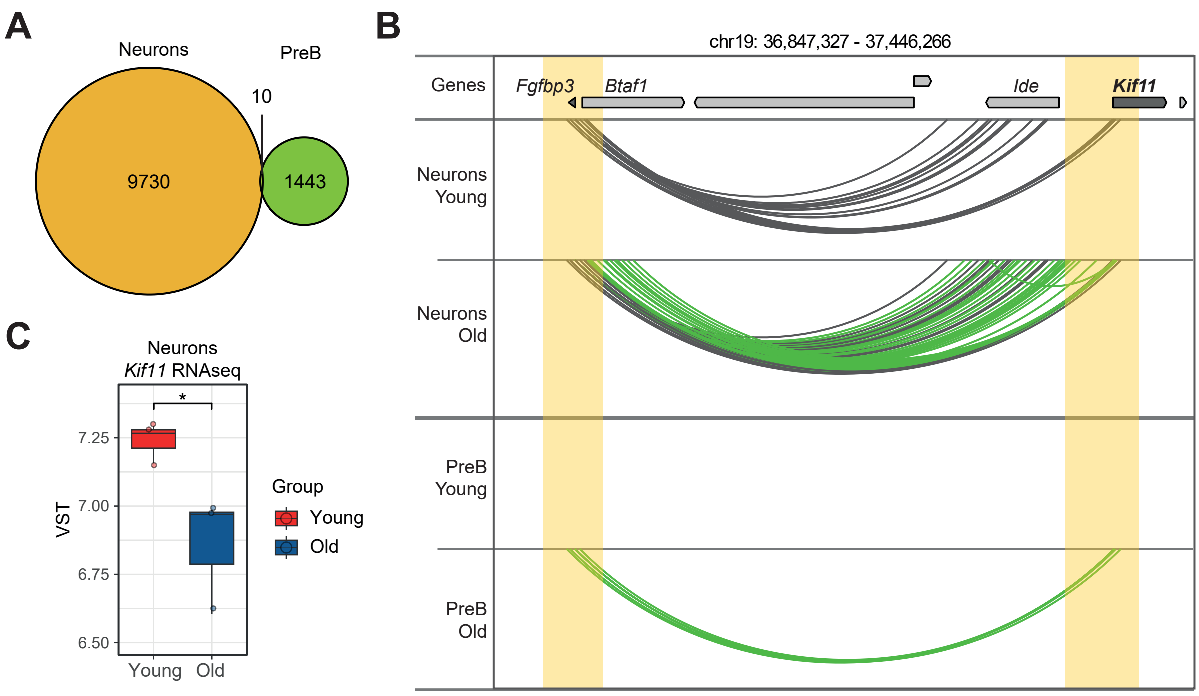


**Figure S4. Neurons and pre-B cells comparison.**

**A.** Venn diagram depicting the number of significant interactions (p-value < 0.05) for aging (old vs young) in neurons (orange) and in pre-B cells (green). The number of overlapping interactions (significant for neurons and pre-B) is highlighted. **B**. *Kif11* and *Fgfbp3* promoters-centered interactions in young and old conditions according to liCHi-C data of neurons and PCHi-C data of pre-B cells. Arcs represent CHiCAGO significant contacts, green arcs highlight those contacts that are found in old mice but not in young mice. Yellow shades depict regions with significantly altered interactions between young and old conditions of both cell types (Table S7). Arrows symbolize gene placement and orientation along the genomic window. **C**. Boxplots indicating the hippocampus RNA-seq expression measurements (VST normalized data) for the *Kif11* gene across young and old mice. Dots denote individual mouse. *p-value < 0.05.

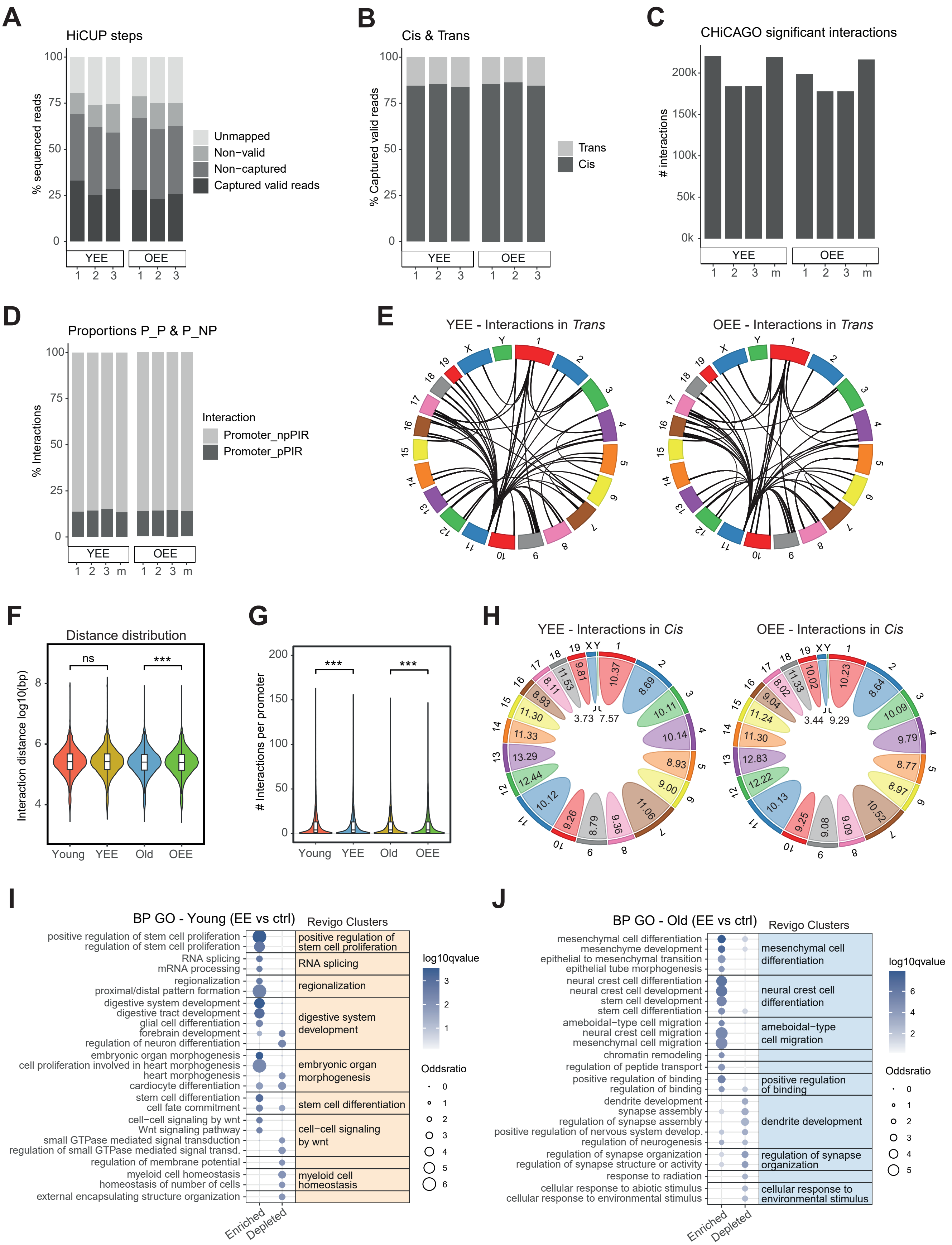
A
HiCUP steps
% sequenced reads
Unmapped
Non−valid
Non−captured
Captured valid reads
YEE
OEE
B
Cis & Trans
% Captured valid reads
Trans
Cis
C
CHiCAGO significant interactions
### interactions
D
Proportions P_P & P_NP
% Interactions
Interaction
Promoter_npPIR
Promoter_pPIR
E
YEE - Interactions in Trans
OEE - Interactions in Trans
F
Distance distribution
Interaction distance log10(bp)
ns
***
Young
YEE
Old
OEE
G
### Interactions per promoter
H
YEE - Interactions in Cis
OEE - Interactions in Cis
I
BP GO - Young (EE vs ctrl)
Revigo Clusters
positive regulation of stem cell proliferation
regulation of stem cell proliferation
RNA splicing
mRNA processing
regionalization
proximal/distal pattern formation
digestive system development
digestive tract development
glial cell differentiation
forebrain development
regulation of neuron differentiation
embryonic organ morphogenesis
cell proliferation involved in heart morphogenesis
heart morphogenesis
cardiocyte differentiation
stem cell differentiation
cell fate commitment
cell−cell signaling by wnt
Wnt signaling pathway
small GTPase mediated signal transduction
regulation of small GTPase mediated signal transd.
regulation of membrane potential
myeloid cell homeostasis
homeostasis of number of cells
external encapsulating structure organization
positive regulation of stem cell proliferation
RNA splicing
regionalization
digestive system development
embryonic organ morphogenesis
stem cell differentiation
cell−cell signaling by wnt
myeloid cell homeostasis
log10qvalue
Oddsratio
Enriched
Depleted
J
BP GO - Old (EE vs ctrl)
Revigo Clusters
mesenchymal cell differentiation
mesenchyme development
epithelial to mesenchymal transition
epithelial tube morphogenesis
neural crest cell differentiation
neural crest cell development
stem cell development
stem cell differentiation
ameboidal−type cell migration
neural crest cell migration
mesenchymal cell migration
chromatin remodeling
regulation of peptide transport
positive regulation of binding
regulation of binding
dendrite development
synapse assembly
regulation of synapse assembly
positive regulation of nervous system develop.
regulation of neurogenesis
regulation of synapse organization
regulation of synapse structure or activity
response to radiation
cellular response to abiotic stimulus
cellular response to environmental stimulus
mesenchymal cell differentiation
neural crest cell differentiation
ameboidal−type cell migration
positive regulation of binding
dendrite development
regulation of synapse organization
cellular response to environmental stimulus
log10qvalue
Oddsratio
Enriched
Depleted

**Figure S5. liCHi-C processing and analysis for EE conditions.**

**A**. Stacked barplots representing proportion of reads passed through the different steps of HiCUP in young with environmental enrichment (YEE) and old with environmental enrichment (OEE) liCHi-C replicates. **B**. Stacked barplots illustrating the cis-trans interactions proportion (in relative percentage) of captured valid reads in YEE and OEE liCHi-C replicates. **C**. Barplots showing the total number of CHiCAGO significant interactions (score > 5) of YEE and OEE liCHi-C replicates and merged samples (m). 1000 (k). **D**. Stacked barplots indicating the proportion of promoter-npPIR significant interactions (promoter and non-promoter PIR) and promoter-pPIR significant interactions (promoter and promoter PIR) of YEE and OEE liCHi-C replicates and merged samples (m). PIR: Promoter Interacting Region. **E**. Circle plots representing *trans* interaction between each mice chromosome for YEE mice (left) and OEE mice (right). **F**. Violin plots showing the distance distribution ($Log_{10}$ base pairs) of CHiCAGO significant interactions of merged samples for YEE and OEE conditions. Wilcoxon rank-sum test was used to compare young and YEE distributions (p-value = 0.602), as well as old and OEE distributions (p-value = 4.495e-10). $^{ns}$p-value > 0.05, ***p-value < 0.001. **G**. Violin plots showing the number of interactions per promoter. Merged samples for YEE and OEE conditions were compared with their respective control conditions, using Wilcoxon signed-rank test for paired samples (p-value: < 2.2e-16 for YEE vs young; 3.698e-13 for OEE vs old). ***p-value < 0.001. **H.** Chord plots representing the *cis* interaction of each mice chromosome (outside number) for YEE mice (left) and OEE mice (right). The width of each chord represents the proportion of *cis* interaction of each chromosome, and the inside numbers indicate the number of *cis* interactions per captured promoter in each chromosome. **I, J.** Bubble plots represent Gene Ontology (GO) terms enrichment based on all Biological Processes (BP) for promoter genes with significantly enriched and depleted interactions (p-value < 0.05) with EE in young mice (**I**) and EE in old mice (**J**). Bubble color intensity represents the statistical significance ($-Log_{10}$ q-value) and dot size reflects odds ratio. Top significant GO terms were grouped following REVIGO clustering, which are named with the most representative term of the group.

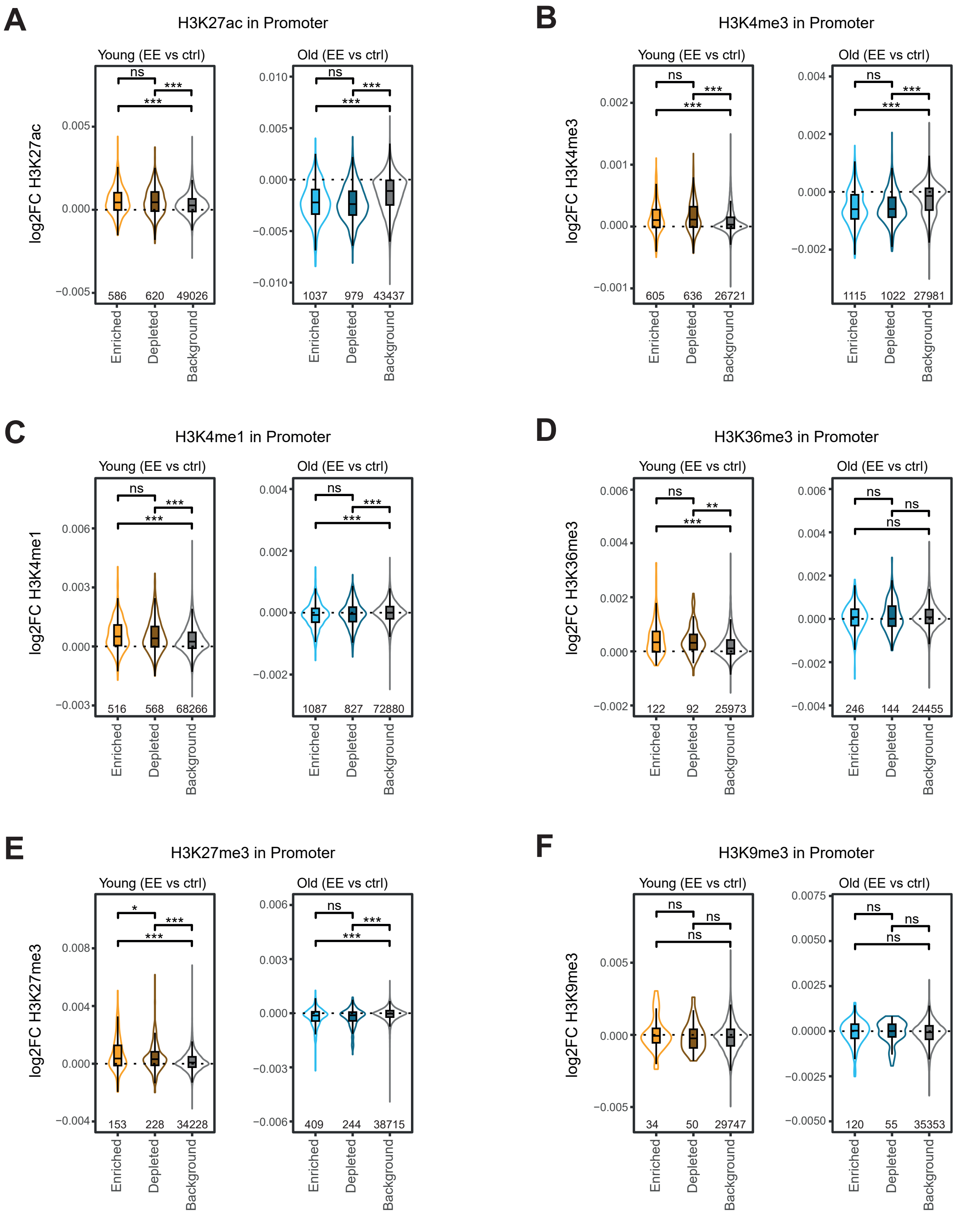


**Figure S6. Histone marks peaks of promoter regions with altered interactions upon EE.**

Violin plots showing the differential (Log$_2$ Fold Change) at promoter regions of histone marks from a previous study[47]: H3K27ac (**A**), H3K4me3 (**B**), H3K4me1 (**C**), H3K36me3 (**D**), H3K27me3 (**E**), H3K9me3 (**F**). Histone mark peaks were analyzed for EE in young and EE in old mice, differentiating between peaks associated with significantly enriched interactions (p-value < 0.05), with significantly depleted interactions (p-value < 0.05) and with non-significantly altered interactions (p-value > 0.05; background). Sample sizes are shown inside the plots. Wilcoxon rank-sum test was used to test whether the distribution histone marks log$_2$FC differed significantly between groups. $^{ns}$p-value > 0.05, *p-value < 0.05, **p-value < 0.01, ***p-value < 0.001.

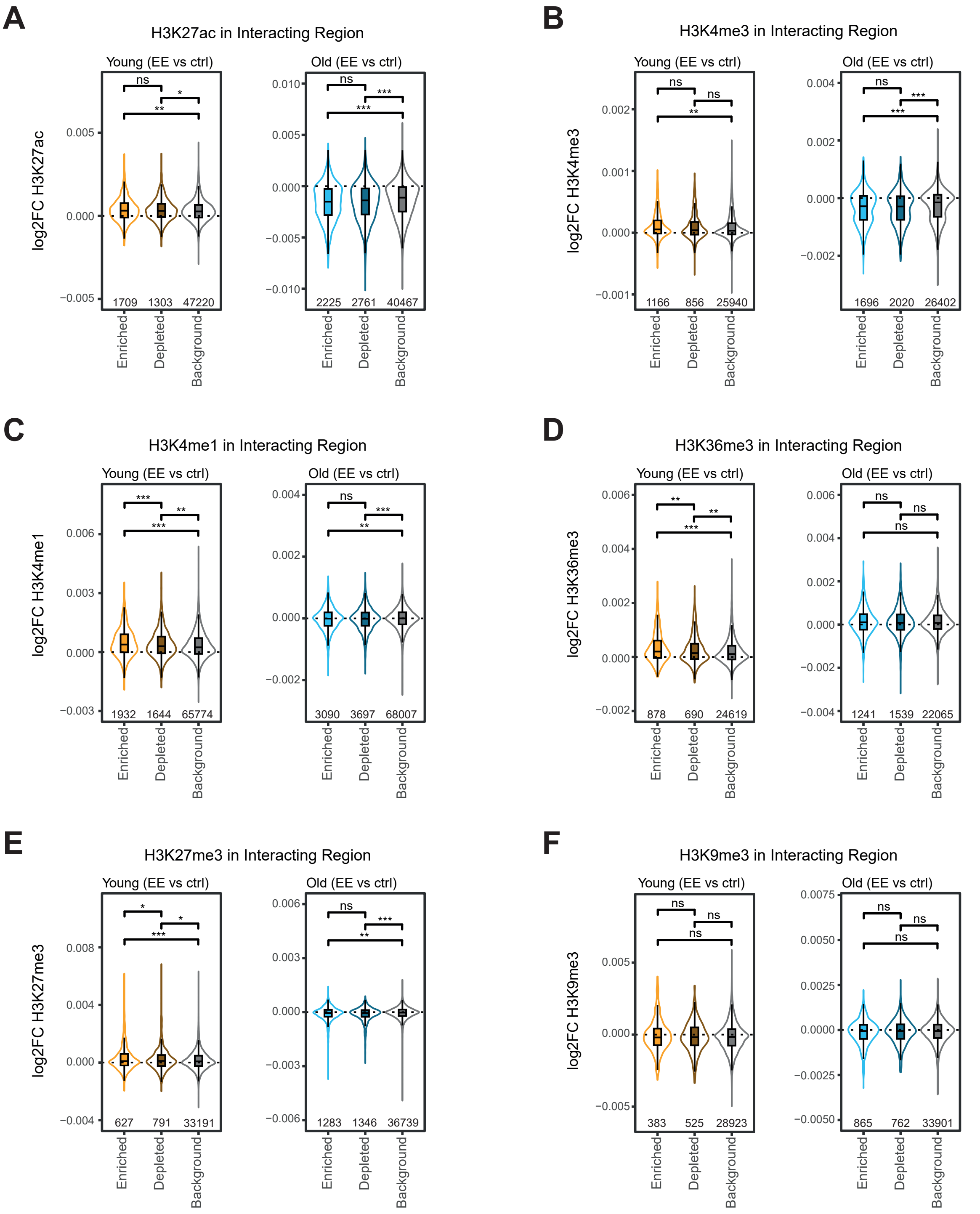


**Figure S7. Histone marks peaks of interacting regions with altered interactions upon EE.**

Violin plots showing the differential ($Log_2$ Fold Change) at interacting regions of histone marks from a previous study[47]: H3K27ac (**A**), H3K4me3 (**B**), H3K4me1 (**C**), H3K36me3 (**D**), H3K27me3 (**E**), H3K9me3 (**F**). Histone mark peaks were analyzed for EE in young and EE in old mice, differentiating between peaks associated with significantly enriched interactions (p-value < 0.05), with significantly depleted interactions (p-value < 0.05) and with non-significantly altered interactions (p-value > 0.05; background). Sample sizes are shown inside the plots. Wilcoxon rank-sum test was used to test whether the distribution histone marks $log_2FC$ differed significantly between groups. $^{ns}$p-value > 0.05, *p-value < 0.05, **p-value < 0.01, ***p-value < 0.001.

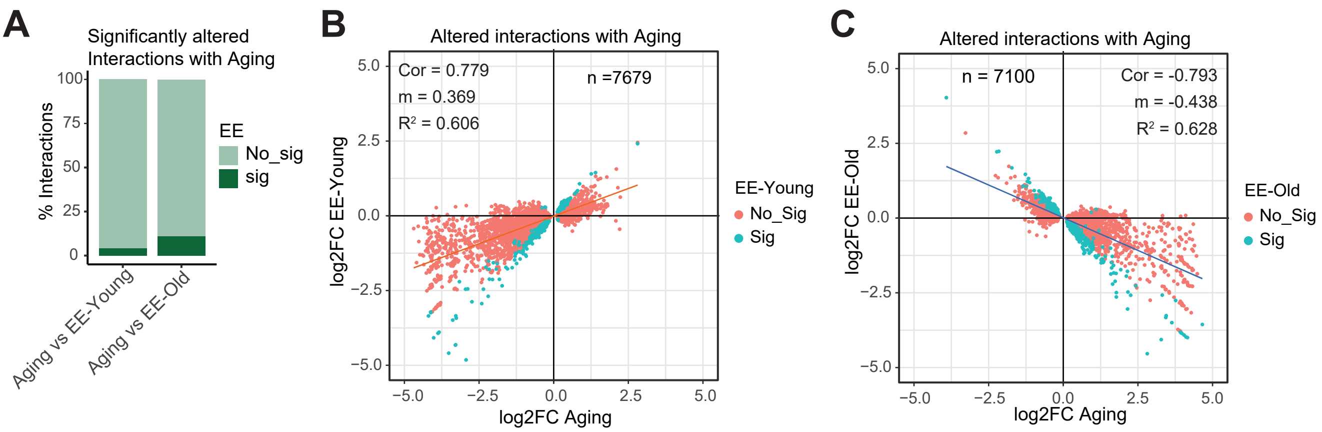


**Figure S8. EE effect on altered interactions with aging.**

**A.** Stacked bar plots illustrating the relative frequency of significantly altered interactions (p-value > 0.05) with aging (old vs young), in relation to its statistical significance with EE in young or in old mice. Dark green corresponds to significant interactions with EE (p-value < 0.05; sig; 4.219% and 10.817% respectively) and light green corresponds to non-significant interactions with EE (p-value > 0.05; No sig). **B.** Scatter plot showing $Log_2$ Fold Change with aging (old vs young) and EE in young mice, of those interactions that are significantly altered at least with aging (p-value < 0.05) (Pearson's correlation score = 0.779; p-value < 2.2e-16). Blue dots highlight interactions that also have a p-value < 0.05 for EE in young mice (Sig), while red dots indicate interactions that are not significant for EE in young mice (No Sig). Orange line indicates the regression line (slope and R-squared value are shown). **C.** Scatter plot representing $Log_2$ Fold Change with aging and EE in old mice, of those interactions that are significantly altered at least with aging (p-value < 0.05) (Pearson's correlation score = -0.793; p-value < 2.2e-16). Blue dots highlight interactions that also have a p-value < 0.05 for EE in old mice (Sig), while red dots indicate interactions that are not significant for EE in old mice (No Sig). Blue line indicates the regression line (slope and R-squared value are shown).

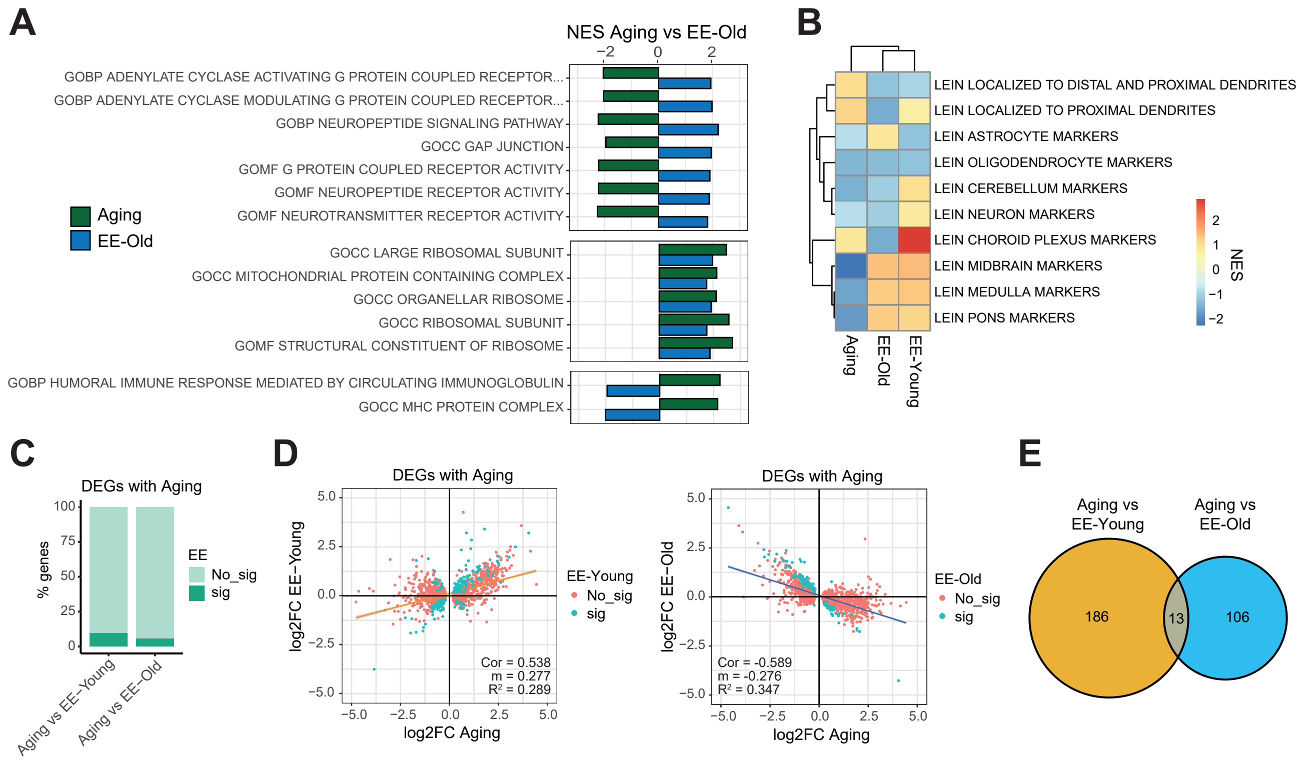


**Figure S9. Gene expression for aging and EE.**

**A**. Barplots reflecting the Normalized Enrichment Score (NES) of GSEA gene sets from M5 collection (MSigDB; Gene Ontologies). Gene sets NES for aging (FDR q-value < 0.05) and EE in old mice (FDR q-value < 0.2) are shown. Gene sets were grouped in clusters based on NES signs. **B**. Heatmap showing the GSEA enrichment of the studied processes (aging, EE in young mice and EE in old mice) towards delimited brain areas denoted by the MSigDB LEIN markers datasets. Color scale indicates the normalized enrichment scores (NES) for each condition. **C**. Stacked bar plots illustrating the relative frequency of DEGs (p-value > 0.05) with aging (old vs young), in relation to its statistical significance with EE in young or in old mice. Dark green corresponds to significant interactions with EE (p-value < 0.05; sig; 9.531% and 5.767% respectively) and light green corresponds to non-significant interactions with EE (p-value > 0.05; No sig). **D.** Scatter plots showing Log$_2$ Fold Change with aging (old vs young) and EE in young mice (left), or Log$_2$ Fold Change with aging and EE in old mice (right), of those interactions that are significantly altered at least with aging (p-value < 0.05) (Pearson's correlation score = 0.538 and -0.589 respectively; p-value < 2.2e-16 in both). Blue dots highlight interactions that also have a p-value < 0.05 for EE (Sig), while red dots indicate interactions that are not significant for EE (No Sig). Orange and blue lines indicate regression lines (slopes and R-squared values are shown). **E.** Venn diagram depicting the number of significant interactions (p-value < 0.05) for aging and EE in young mice (orange), or for aging and EE in old mice (light blue). The number of overlapping interactions (significant for aging and both types of EE) is highlighted in grey. The overlap has a representation factor of 10.2 (p-value = 4.366 x $10^{-10}$; universe = 18715 genes) using hypergeometric distribution.

#### SUPPLEMENTARY TABLE LEGENDS

**Table S1**. liCHi-C analysis. Read counts passed through the different steps of HiCUP and CHiCAGO, for young, old, YEE and OEE replicates are shown, as well as CHiCAGO values for merged samples.

**Table S2**. Significant enrichments in GO terms (q-value < 0.05) for promoters with significantly altered interactions (enriched or depleted), considering Biological processes (All BP) or Nervous system-associated biological processes (Nervous BP). All GO terms for Aging, EE-Young and EE-Old are shown. GO enrichments for the Aging-EE comparatives (Aging vs EE-Young; Aging vs EE-Old) are also included.

**Table S3**. Mapping details for RNA-seq replicates.

**Table S4**. Enrichments in GSEA gene sets, considering an FDR q-value threshold of 0.05 for Aging and 0.2 for EE. Gene sets enrichments for every comparative (EE-Young vs EE-Old; Aging vs EE-Young; Aging vs EE-Old) are shown. Also, Aging vs EE-Old comparative for M5 collection, and enrichments with LEIN markers, are included. All enrichment score (ES) and normalized enrichment score (NES) are indicated.

**Table S5**. Integration of liCHi-C and RNA-seq data (p-value < 0.05). Fold change ($Log_2$ Fold Change) and statistical significance (p-value) of differentially expressed genes (DEGs) with altered interactions in Aging, EE-Young and EE-Old, are shown.

**Table S6**. Chicdiff significant altered interactions (p-value < 0.05) for example gene promoters: *Selenow*, *Htr7*, *Sema5b*, *Ndnf*, *Fosl2*, *Grm4*.

**Table S7**. Chicdiff significant altered interactions (p-value < 0.05) overlapping between neurons and pre-B cells.

**Table S8**. Integration of liCHi-C data (p-value < 0.05) and RNA-seq data (p-value < 0.2), comparing between Aging and EE (Aging vs EE-Young, and Aging vs EE-Old). Fold change ($Log_2$ Fold Change) and statistical significance (p-value) of genes with altered interactions are shown.

**Table S9.** Primers and adapters for 3C quality controls and liCHi-C library construction.

**Table S10.** Nervous System-associated Biological Processes (BP) Gene Ontologies (GO) terms. The source subsets of each GO terms are indicated: Neural-Immune Gene Ontology (NIGO); Synaptic Gene Ontologies (SynGO).
